# An Epigenetic Priming Mechanism Mediated by Nutrient Sensing Regulates Transcriptional Output

**DOI:** 10.1101/2020.09.01.278127

**Authors:** Natalia Stec, Katja Doerfel, Kelly Hills-Muckey, Victoria M. Ettorre, Sevinc Ercan, Wolfgang Keil, C. M. Hammell

## Abstract

While precise tuning of gene expression levels is critical for most developmental pathways, the mechanisms by which the transcriptional output of dosage-sensitive molecules is established or modulated by the environment remain poorly understood. Here, we provide a mechanistic framework for how the conserved transcription factor BLMP-1/Blimp1 operates as a pioneer factor to decompact chromatin near its target loci hours before transcriptional activation and by doing so, regulates both the duration and amplitude of subsequent target gene transcription. This priming mechanism is genetically separable from the mechanisms that establish the timing of transcriptional induction and functions to canalize aspects of cell-fate specification, animal size regulation, and molting. A key feature of the BLMP-1-dependent transcriptional priming mechanism is that chromatin decompaction is initially established during embryogenesis and maintained throughout larval development by nutrient sensing. This anticipatory mechanism integrates transcriptional output with environmental conditions and is essential for resuming normal temporal patterning after animals exit nutrient-mediated developmental arrests.

## INTRODUCTION

Many developmental systems produce invariable phenotypes despite being confronted with perturbations, such as fluctuations in temperature and nutrient availability. How this robustness emerges from developmental gene regulatory networks (GRNs) and how aspects of precision and flexibility are balanced to generate robust developmental programs is a major open question in developmental biology. *C. elegans* serves as a unique genetic model for unraveling the mechanisms underlying precision and robustness of gene regulation during development. This is in part because the GRNs that control the essentially invariant cell lineage can generate the identical developmental outcomes in the context of diverse stochastic, environmental or genetic perturbations. A striking example of such a GRN is the so-called heterochronic pathway which controls the timing, sequence and synchrony of stage-specific patterns of cell divisions throughout post-embryonic development (Ambros and Horvitz, 1984). Analysis of this GRN indicates that it generates modular patterns of gene expression that occur in the context of the repetitive post-embryonic molting cycle (Ambros and Horvitz, 1984) which, in turn, demarcates the four successive larval stages (L1-L4). Transitions from one temporal program to the next occur through the expression of distinct microRNAs (miRNAs) that down-regulate the expression of their temporal selector gene targets in a strict dosage-sensitive manner. Ectopic or abnormally lower or higher doses of heterochronic genes (e.g. miRNAs) result in the wholesale skipping or reiteration of stage-specific developmental programs (Rougvie and Moss, 2013, Feinbaum and Ambros, 1999; Li et al., 2005; Perales et al., 2014; Van Wynsberghe and Pasquinelli, 2014). While the regulatory interactions that establish the sequence of stage-specific events are known, we have very little understanding of how the precise timing or levels of gene expression within this GRN are established and how both of these are buffered against perturbations to achieve robust temporal development of the animal.

High-temporal resolution transcriptomics studies have outlined a number of remarkable features associated with global gene expression patterns in developing *C. elegans* larva (Hendriks et al., 2014; Kim et al., 2013). These studies indicate that between 10 to 20% of the post-embryonic transcriptome exhibits highly reproducible periodic expression patterns. Periodic transcription occurs in a variety of environmental conditions and is independent of life history, indicating that it is under tight genetic control. Importantly, the transcriptional rhythm follows the cycle of post-embryonic molting, a process that demarcates patterns of stage-specific developmental programs. Under various environmental conditions that modulate overall developmental pace, the timing of transcription onset scales accordingly, such that the phase of transcription onset within the molting cycle is preserved (Hendriks et al., 2014; Kim et al., 2013). Currently, it is not known how these transcriptional rhythms are generated, how they are integrated into the execution of stage-specific cellular programs, and how environmental or internal cues modulate features of these transcriptional patterns to achieve robust progression through developmental programs, even after prolonged developmental arrests.

The heterochronic GRN is integrated with global aspects of transcription as each of the miRNAs in this pathway exhibits an oscillatory expression. The most promising candidate gene that integrates the rhythm of *C. elegans* post-embryonic molting to changes in repetative transcriptional patterns is *lin-42*. The *lin-42* gene encodes the *C. elegans* ortholog of PERIOD/Per proteins that are an essential component of the circadian clock in mice, *Drosophila* and humans (Hurley et al., 2016; Jeon et al., 1999; Monsalve et al., 2011). Similar to PERIOD, LIN-42 functions as a transcriptional repressor and exhibits a cyclical expression pattern In contrast to the temperature-invariant approximate 24-hour periodicity of *Period, lin-42* transcription is tied to the larval molting cycle, and thus, not temperature compensated (Monsalve et al., 2011; Perales et al. 2014). Hypomorphic mutations in *lin-42* cause animals to prematurely execute adult-specific gene expression patterns in the L4 stage of larval development and these phenotypes are caused by the cumulative, premature over-expression of multiple heterochronic miRNAs (McCulloch and Rougvie, 2014; Perales et al., 2014; Van Wynsberghe and Pasquinelli, 2014). In addition to these temporal patterning phenotypes, *lin-42* mutants also exhibit lengthened and irregular molting cycles (Monsalve et al. 2011; Tennessen et al., 2010). The mechanisms by which LIN-42 negatively regulates heterochronic miRNA transcriptional output remain unknown. Furthermore, it is not known if other regulatory components exist that antagonize LIN-42 functions to ensure normal transcriptional regulation of developmental timing genes.

Here, we leverage the developmental phenotypes of *lin-42* mutations to identify BLMP-1 and ELT-3 as conserved transcription factors (TFs) that modulate the transcriptional output of many cyclically expressed genes to provide developmental robustness. We demonstrate that these TFs coordinate a plethora of post-embryonic developmental programs and we present a unique role for BLMP-1 in transcriptional priming that involves chromatin decompaction of target loci. Specifically, we show that BLMP-1 activity modulates the duration and amplitude of transcription of its targets without altering the timing of expression within the repetitive transcriptional cycle. We hypothesize that BLMP-1 functions specifically to assure proper transcriptional output in diverse environmental conditions. Consistent with this model, we demonstrate that the BLMP-1-dependent remodeling of chromatin is regulated by nutritional inputs and becomes essential when animals resume development after nutrient-mediated developmental arrests.

## RESULTS

### *blmp-1* and *elt-3* Function in a partially redundant manner to control temporal patterning during larval development

To identify genes that antagonize the function of LIN-42 to modulate transcriptional output, we took advantage of a highly penetrant phenotype of the *lin-42* loss of function allele *lin-42(n0189)* – the precocious expression of an adult-specific *col-19::GFP* reporter in both seam and syncytial hyp7 cells of the hypodermis (Figure 1A) (Abrahante et al., 1998). We used bacterial-mediated RNAi to deplete the expression of individual *C. elegans* TFs in *lin-42(lf)* animals and then scored *col-19::GFP* expression in F_1_ progeny. The most reproducible suppressors of *lin-42* precocious phenotypes were bacterial RNAi clones expressing dsRNAs that target the *blmp-1* and *elt-3* loci (data not shown). We validated these findings with null alleles of these genes (*blmp-1(0)* and *elt-3(0)*) which revealed a redundancy between *blmp-1* and *elt-3* that is distributed differentially across hypodermal cell types (Figure 1 A and B). Specifically, removal of *blmp-1* activity in *lin-42(lf)* mutants completely suppresses the precocious *col-19::GFP* expression in the lateral seam cells of L4-staged animals and only partially prevents early expression in hyp7 cells (Figure 1A and B). *lin-42(lf); elt-3(0)* mutants were only partially suppressed for the precocious *col-19::GFP* expression phenotypes. In contrast, combining *blmp-1(0)* and *elt-3(0)* mutations with *lin-42(lf)* mutations completely alleviated precocious *col-19::GFP* expression in both the seam and hyp7 cells of a majority of L4-staged animals (Figure 1A and B). This cell-type specific redundancy is likely due to the partially overlapping expression patterns of these TFs in hypodermal cells throughout larval development and indicate that BLMP-1 and ELT-3 are required for *lin-42(lf)* phenotypes(Cao et al., 2017) (Figure 1C).

**Figure 1.**
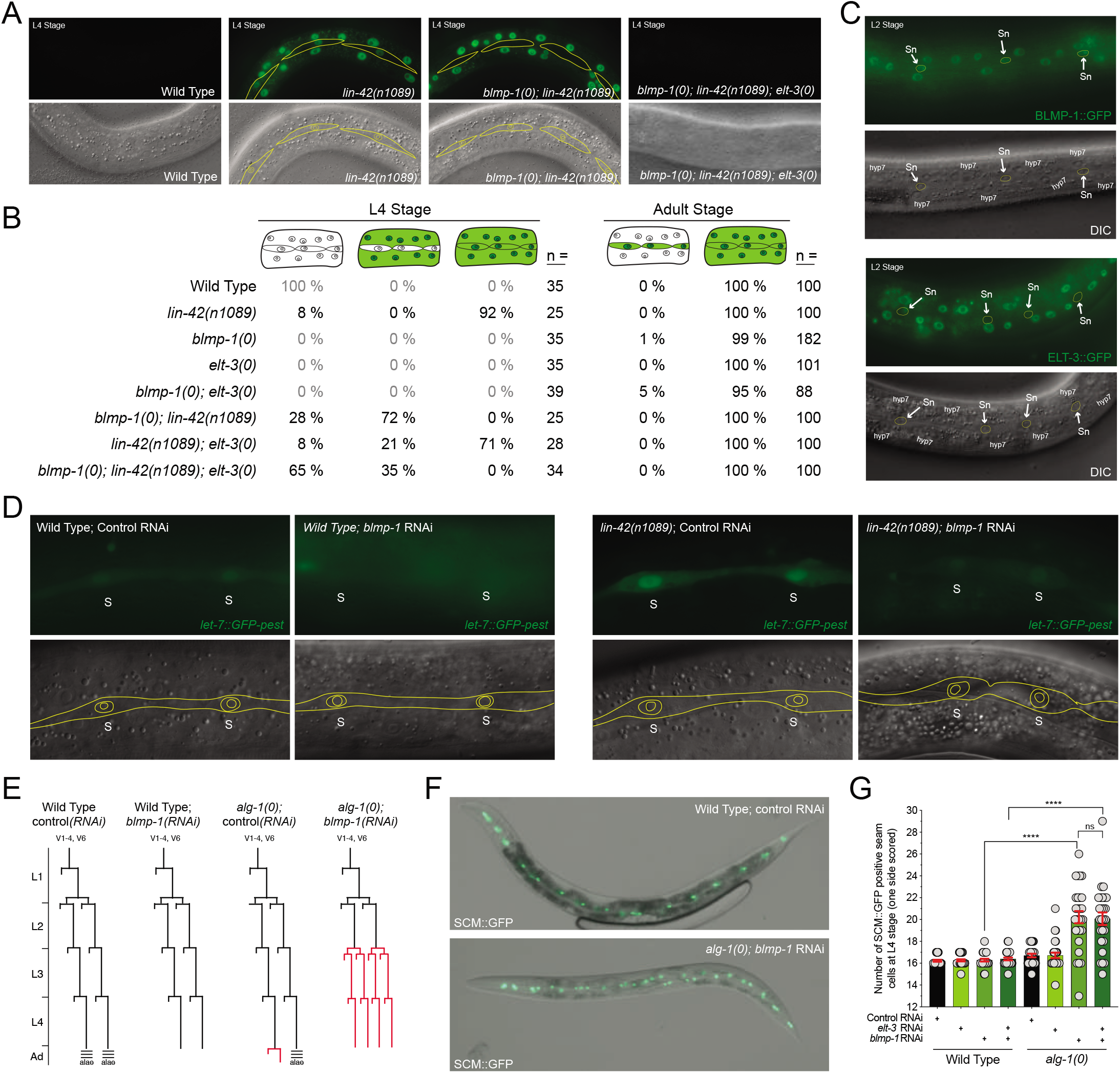
*blmp-1* and *elt-3* antagonize *lin-42* activity to control the expression of heterochronic miRNAs. **(A)** Representative *col-19::GFP* expression patterns in wild-type, *lin-42(lf), blmp-1(0); lin-42(lf)*, and *blmp-1(0); lin-42(lf); elt-3(0)* animals. **(B)** A quantification of the *col-19::GFP* expression phenotypes of various mutant combinations in L4- and adult-staged animals. **(C)** Expression patterns of BLMP-1::GFP and ELT-3::GFP transgenes in hypodermal cells. Sn = Seam cell nuclei. **(D)** Representative images of mid-L4-staged wild-type or *lin-42(lf)* animals expressing a *let-7::GFP-pest* transcriptional reporter. S = seam cell. **(E)** Lateral seam cell lineage of wild-type, *blmp-1*, and *alg-1* mutants and experiments outlined in panel F. **(F)** SCM::GFP is expressed in lateral seam cells and can be used to monitor cell division patterns during larval development. Mid-L4-staged *alg-1(0); blmp-1(RNAi)* animals exhibit a supernumerary number of SCM::GFP seam cells indicative of an inappropriate reiteration of seam cell division programs. **(G)** Quantification of lateral seam cell numbers of mid-L4-staged animals exposed to bacteria expressing dsRNAs complementary to *elt-3, blmp-1* or both *blmp-1* and *elt-3*. Asterisks indicate statistically significant differences in phenotype (p = < 0.0001) calculated from a two-tailed chi-square analysis.

Because many of the developmental phenotypes associated with *lin-42* mutations are due to elevated expression of heterochronic miRNAs, we sought to determine if the genetic relationships between *blmp-1, elt-3 a*nd *lin-42* are mediated at the level of transcription. A *let-7::GFP-pest* reporter is expressed in the lateral seam cells of L3 and L4-staged wild-type animals in a pulsatile fashion (once per stage). *lin-42(lf)* animals maintain pulsatile expression patterns but the amplitude of *let-7::GFP-pest* expression is elevated in lateral seam cells (Perales et al. 2014). Consistent with *blmp-1* functioning up-stream of *lin-42* to control *let-7* transcriptional output, *let-7::GFP-pest* expression in *lin-42(lf); blmp-1(RNAi)* animals was reduced to almost wild-type expression levels. Furthermore, *let-7::GFP-pest* expression in wild-type *blmp-1-(RNAi)* animals is reduced compared to control RNAi conditions consistent with a normal function of BLMP-1 in modulating transcriptional output of this miRNA. RNAi depletion of *elt-3* expression did not alter *let-7::GFP* expression in L4-staged wild-type or *lin-42(lf)* animals (data not shown).

Previous analysis of *blmp-1(0)* mutations indicate that BLMP-1 plays an important role in terminal cell fate specification of the hypodermis (Horn et al., 2014). We sought to determine if *blmp-1* functions earlier in temporal cell fate specification when cells are proliferating. To this end, we assayed genetic interactions between *blmp-1* and *alg-1*, the latter encoding one of the two core miRISC argonaute components. *alg-1(0)* mutants ineffectively process most miRNAs (including heterochronic microRNAs) and exhibit relatively mild temporal patterning defects that resemble those associated with hypomorphic alleles of *let-7* (Hammell et al. 2009). To assay interactions, we quantified the number of seam cells at the mid L4 stage in wild-type or *alg-1(0)* mutant animals in conditions where we reduced *blmp-1* and/or *elt-3* activity using RNAi. In wild-type animals, lateral seam cells exhibit highly reproducible cell division patterns that occur at specific stages of larval development (Figure 1E) (Sulston and Horvitz, 1977). Defects in this lineage can be monitored by counting seam cell numbers in the L4 larval stage. Depletion of *elt-3* or *blmp-1* in wild-type animals resulted in the normal number of seam cells in the L4 stage. We observed a significant increase in the number of SCM::GFP(+) seam cells in *alg-1(0); blmp-1* RNAi animals (Figure 1F and G). This expansion of lateral seam cell number resembles an inappropriate reiteration of earlier, L2-specific proliferative cell division programs (Figure 1E). Depletion of *blmp-1* in other mildly reiterative heterochronic mutants also resulted in similar enhancement of cell lineage and *col-19::GFP* expression phenotypes (Figure S1). *elt-3* RNAi did not significantly alter lateral seam cell numbers in *alg-1(0)* animals or enhance *alg-1(0); blmp-1(RNAi)* phenotypes. This indicates that BLMP-1 functions early in larval development to control temporal cell fates in the hypodermis.

### *blmp-1* and *elt-3* regulate multiple dosage-dependent developmental processes

Development of *C. elegans* larva results in a fixed number of cells and a stereotypical size under standard growth conditions. Mutations in a number of pathways (including nutrient signaling, TGFβ and MAP kinase pathways) or in structural components of the cuticle alter organismal size (Tuck, 2014). A shared feature of these regulatory and structural systems is that they function in a dose-sensitive manner wherein changes in the expression levels of individual elements within these pathways or components result in changes in animal morphology. As shown in Figure 2A, *blmp-1(0); elt-3(0)* animals exhibit a severe dumpy (*dpy*) phenotype. The *dpy* phenotype of *blmp-1(0); elt-3(0)* mutants manifests during post-embryonic development as freshly hatched *blmp-1(0); elt-3(0)* larvae are indistinguishable from wild-type animals (data not shown). *blmp-1(0)* single mutants exhibit less severe alterations in size and *elt-3(0)* animals are structurally normal (Figure 2A and B). These results demonstrate that in addition to modulating aspects of temporal cell-fate specification, *blmp-1* and *elt-3* cooperate to control animal morphology likely through the co-regulation of hypodermally expressed targets.

**Figure 2.**
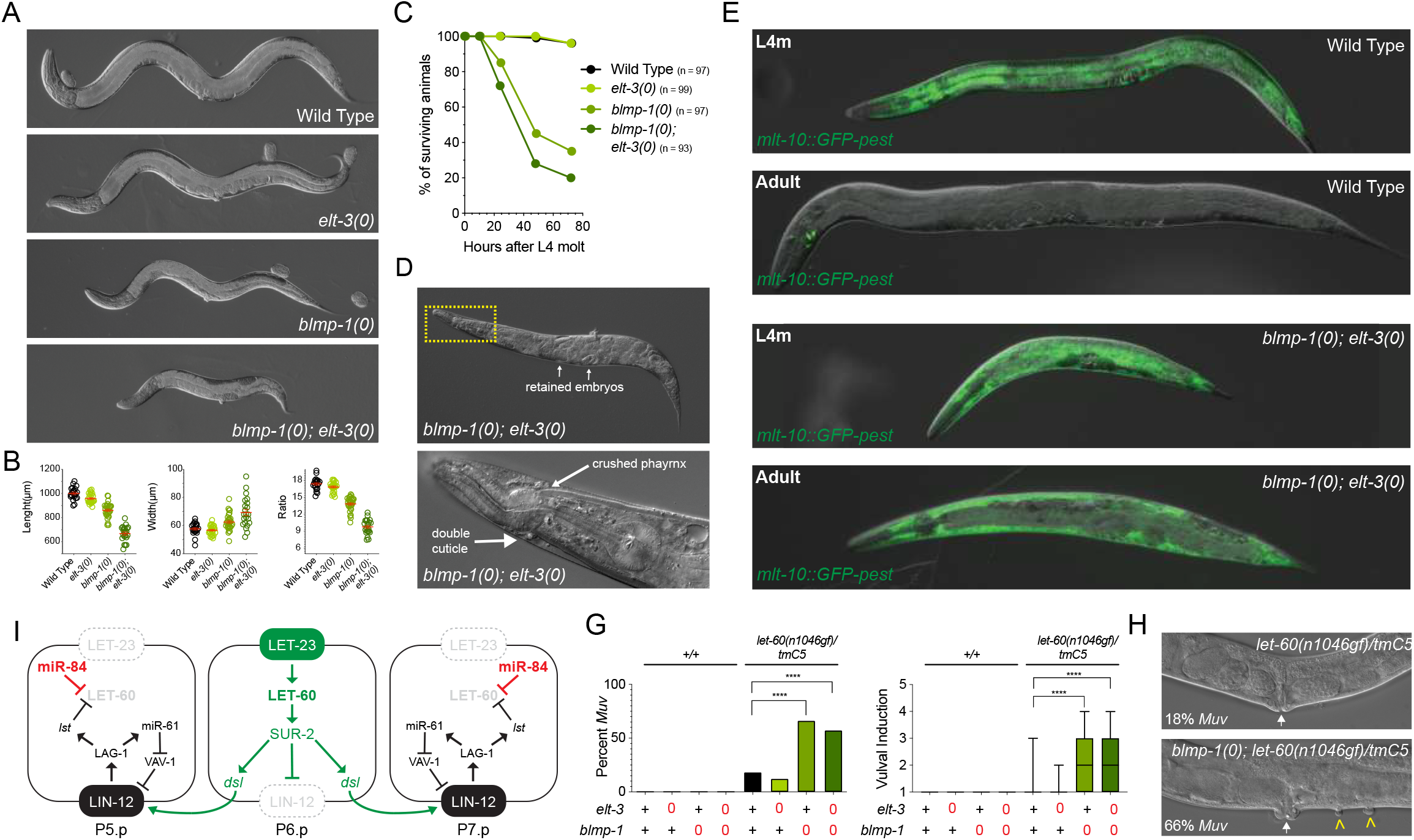
*blmp-1* and *elt-3* regulate several additional morphological, behavioral, and cellular phenotypes. **(A)** *blmp-1(0); elt-3(0)* animals exhibit a synthetic Dpy phenotype that manifests during larval development. **(B)** Quantification of average animal length and width of young adult animals of the indicated genotypes (n = >20). Red bars indicate the mean and SEM. **(C)** *blmp-1(0)* and *blmp-1(0); elt-3(0)* animals prematurely die after the L4-adult transition. **(D)** The lethality of *blmp-1(0); elt-3(0)* adult animals correlates with an inability to shed the cuticle of a supernumerary molt. **(E)** *blmp-1(0); elt-3(0)* inappropriately reanimate the expression of a *mlt-10::GFP-pest* transcriptional reporter during adulthood. Animals were imaged at the L4-adult transition or as gravid adults. **(F)** Major components of the vulval induction pathway (see text for details). Components that are genetically involved in vulval induction are colored green whereas those involved in Notch/lateral inhibition are colored black. The *let-7* family miRNA miR-84 (red) that regulates *let-60* expression is temporally expressed in presumptive 2° vulval cells. **(G)** *blmp-1(0)* mutations enhance the expressivity and penetrance of *let-60(n1046gf) muv* phenotypes. Animals of the indicated genotypes were scored for the number of vulval protrusions present on the ventral side at the young adult stage (n = >90). Brackets indicate statistically significant differences in phenotype calculated from a two-tailed chi-square analysis. **(H)** Representative images depicting Muv phenotypes associated with the indicated genotypes.

A majority of *blmp-1(0); elt-3(0)* animals precociously die two days after they reach adulthood due to the internal hatching of fertilized embryos (Figure 2C). Examination of dying *blmp-1(0); elt-3(0)* animals revealed that most are trapped in a partially shed cuticle (Figure 2D). Lethality is preceded by a reduction in movement and pharyngeal pumping approximately 8-10 hours after the L4-to-adult transition; a phenotype shared with other mutants that fail to post-transcriptionally down-regulate the adult expression of key genes involved in larval molting (Hayes et al., 2006). To determine if *blmp-1(0); elt-3(0)* animals inappropriately re-initiate molting programs during adulthood, we directly monitored the expression of a molting-specific GFP-pest transcriptional reporter, *mlt-10::GFP-pest*, in wild-type, single and double mutant animals.. Wild type animals never expressed *mlt-10::GFP* as adults (n = 100) while a majority of *blmp-1(0); elt-3(0)* animals re-expressed the *mlt-10::GFP-pest* reporter (54%; n = 155) as adults (Figure 2E). *blmp-1(0)* single mutants also express *mlt-10::GFP-pest* within the same time period but with a much lower penetrance (18%, n = 137), consistent with previous observations using *blmp-1(RNAi)* (Frand et al., 2005). These data indicate that *blmp-1* and *elt-3* function redundantly to halt the behavioral and physiological molting programs once animals transition into adulthood.

Development of the vulval structure in the L3 stage requires the integration of a number of interconnected signaling pathways that establish specific cell fates in vulval precursor cells (Figure 2F). One of the key initiating components in this process is *let-60*, encoding the *C. elegans* ortholog of the human Ras oncogene. Continued *let-60* activation commits vulval precursor cells (VPCs) to a 1° vulval cell fate specification pattern (Figure 2F). The induction of this fate specification is sensitive to the level of *let-60* activity (Beitel et al., 1990; Han and Sternberg, 1990). For instance, the semi-dominant *let-60(n1046gf)* allele can induce ectopic vulval protrusions on the ventral surface (multi-vulva; *muv*) in 18% of animals harboring a single copy of this mutation (Figure 2G and H). We used this sensitized genetic context to determine whether *blmp-1(0)* and/or *elt-3(0)* mutations (or the combination of these mutations) leads to an elevation in VPC induction in *let-60(n1046)/+* animals. Lack of *blmp-1* activity significantly increased the penetrance and expressivity of the *muv* phenotype in *let-60(n1046)/+* animals (Figure G and H). *elt-3(0)* mutations did not contribute to *let-60*-dependent phenotypes consistent with a lack of significant ELT-3 expression in these cell types (Figure 3G) (Cao et al., 2017). Together, these results indicate that *blmp-1* and *elt-3* function in a partially redundant manner to regulate a multitude of developmental aspects, from animal morphology, to behavior to cell fate specification.

**Figure 3.**
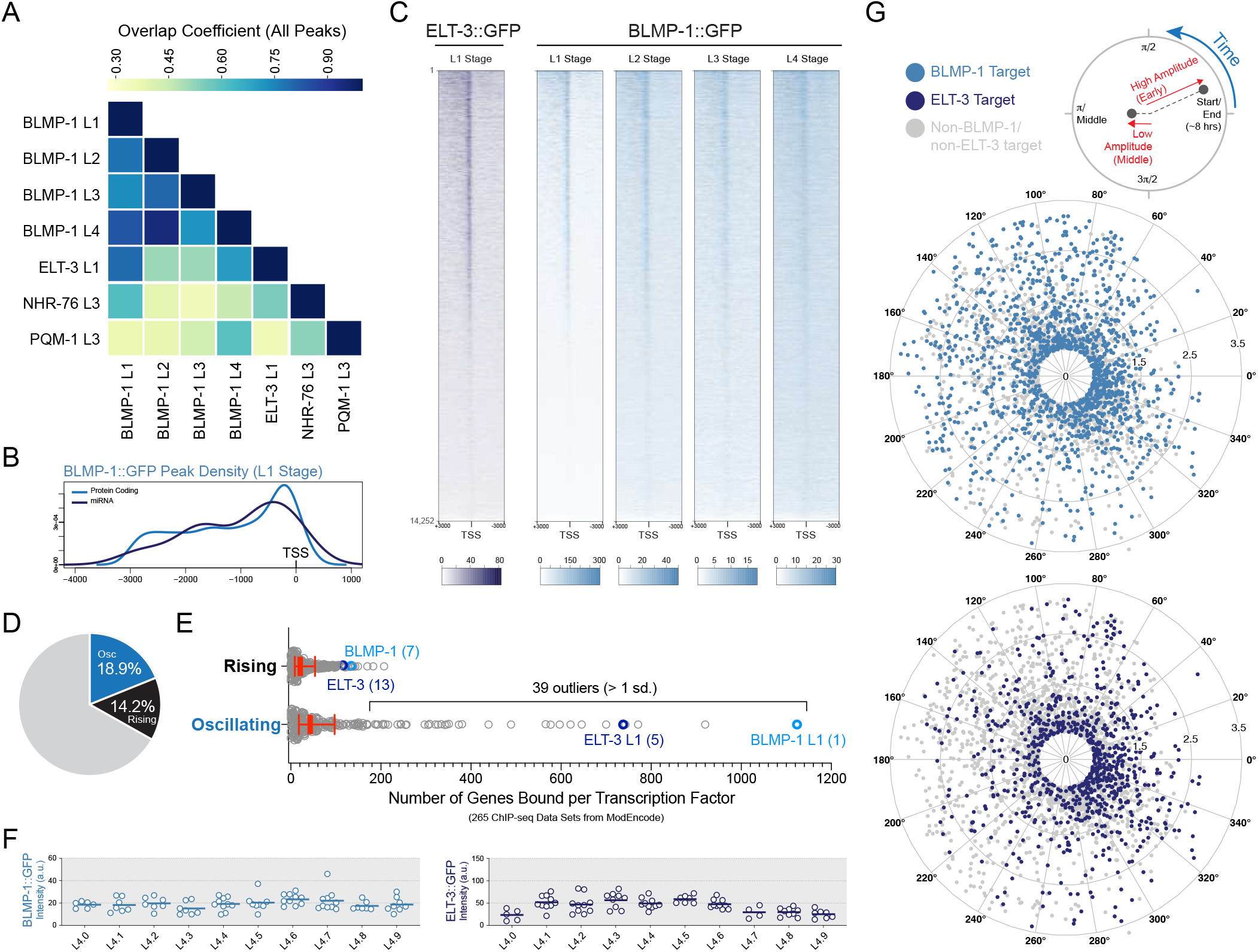
BLMP-1 and ELT-3 bind similar sites in the genome and are associated with the putative regulatory regions of gens that exhibit cyclical expression patterns. **(A)** A matrix of global pairwise factor co-association strengths as defined by promoter interval statistics indicate that BLMP-1 and ELT-3 bind similar genomic regions. **(B)** The distribution of the peak summit positions of BLMP-1 ChIP-seq data mapped relative to the TSS of protein coding genes or to the +1 nucleotide of the DNA sequence encoding the miRNA precursor RNA. **(C)** A heat map of ChIP-seq signals +/- 3 kb centered on the TSS of each of the 14,252 protein coding genes measured in the developmental RNA-seq data set (Hendriks et al., 2014). Genes are organized for each heat map in a 1:1 fashion according to their density of unique reads present in the BLMP-1 L1-stage ChIP-seq sample. **(D)** The *C. elegans* larval transcriptome can be categorized into two dynamic expression classes (rising and oscillating) that are distinct from the predominantly flat expression pattern of most mRNAs (grey region; 66.9%) (Hendriks et al., 2014). **(E)** A graph displaying the number of predicted rising or oscillating class targets for each of the 172 TFs assayed by the ModEncode Projects (data derived from a total of 265 ChIP-seq data sets assayed). Red bars indicate the median (thick) with the interquartile range (thin). **(F)** Quantification of GFP intensity for translational reporters of BLMP-1 and ELT-3 during the L4 stage. **(G)** A radar chart plotting oscillation amplitude over the phase of peak expression reveals that BLMP-1 and ELT-3 bind the putative regulatory regions of periodically expressed genes in all phases of the cyclical transcriptional cycle.

### BLMP-1 and ELT-3 co-target overlapping gene sets

Our observations that *blmp-1* and *elt-3* genetically interact with *lin-42* to control the transcription of cyclically expressed miRNAs (e.g. *let-7*, Figure 1D) and that *blmp-1(0); elt-3(0)* double mutants exhibit pleiotropic defects led us to hypothesize that these TFs may function more broadly to control dynamic gene expression. To define the full repertoire of BLMP-1 and ELT-3 transcriptional targets, we complemented L1-staged BLMP-1 and ELT-3 ChIP-seq data obtained in the modENCODE project with additional ChIP-seq experiments performed in extracts made from L2-, L3-, and L4-staged animals (Araya et al., 2014; Niu et al., 2011). Analysis of these combined data indicates that the regions bound by BLMP-1 in the first larval stage are also bound in subsequent larval stages. Furthermore, the binding sites of BLMP-1 in the L1-L4 stages significantly overlap with regions also bound by ELT-3 in the L1 stage (Figure 3A). A majority of BLMP-1 peaks are enriched within 500bps upstream of the transcriptional start site (TSS) but the enrichment also extends to 5’ intergenic regions with a sharp drop-off approximately 3kb upstream of gene bodies of both protein coding and miRNA genes (Figure 3B and C). Peaks from all datasets were assigned to the nearest proximal gene (both coding and non-coding) using these parameters (Table S1).

To characterize the relationship between BLMP-1, ELT-3 and their targets in more detail, we initially focused our analysis on protein coding genes whose expression can be reliably detected in developing larvae. Our analysis employed multiple, high-resolution, and genome-wide time course analyses of gene expression that parsed the larval developmental transcriptome into distinct temporal expression classes (Hendriks et al., 2014; Kim et al., 2013). Both studies identified significant, overlapping sets of transcripts (between 12% and 18.9% of the transcriptome) that exhibit cyclical expression patterns. Hendriks et al. classified two types of dynamic expression: a first class of transcripts that exhibit an oscillatory or cyclical patterns (osc) that are tied to the molting cycle (osc; 18.9%; 2,718 of 14,378 total genes) or a second class of genes whose expression increases monotonically throughout larval development (rising; 14.2%) (Figure 3D). To compare how BLMP-1 and ELT-3 targets may be distinguished from the targets of other TFs, we analyzed additional publicly available ChIP-seq data sets obtained by the modENCODE project. This data included the targets of 170 additional *C. elegans* TFs that represent diverse classes of TFs (Araya et al., 2014; Niu et al., 2011). Using the same target assignment criteria as above, we found that most TFs are associated with a limited number of target genes that exhibit a rising expression program (median targets bound = 21 per TF; interquartile range = 9 to 54)(Figure 3E; Table S2). This relationship was also conserved for the associations between bulk TFs and the oscillatory class of target genes (median of 43 targets per TF; interquartile range = 18 to 97). Surprisingly, we found that a limited subset of 39 TFs is disproportionally associated with a large fraction of the oscillatory class of genes (Figure 3E). Primary amongst this unique group of 39 TFs were BLMP-1 and ELT-3 (with 1124 and 738 of the 2718 cyclically expressed targets bound per TF, respectively) (Figure 3E). We performed a similar analysis of the Kim et al. RNA-seq data to determine if ELT-3 or BLMP-1 binding sites are also enriched in the promoters of cyclically expressed transcripts as defined by their temporal classifications. Our analysis found that BLMP-1 binding sites, but not ELT-3 binding sites, are enriched in the promoters of each cyclically expressed genes class and de-enriched for those that exhibit monotonic expression patterns (Figure S3 and Table S3).

One robust feature of cyclically expressed mRNAs is that the timing of expression within each larval stage occurs in the same phase of each transcriptional cycle. This expression/phase relationship is preserved even when animals are grown at lower or higher temperatures which lengthens or shortens the periodicity of these patterns, respectively (Hendriks et al., 2014; Kim et al., 2013). Because BLMP-1 and ELT-3 are constitutively expressed through each larval stage (Figure 3F), we aimed to determine if the subset of BLMP-1 and ELT-3 targets that exhibit cyclical expression patterns were concentrated in any particular phase of expression during each larval stage. To compare these patterns, we plotted the peak phase (in degrees/radians of a repetitive cycle) and expression level (log2 change in expression as an increasing radius from the center) for each of the cyclically expressed transcripts on a radial chart (Figure 3G). ELT-3 and BLMP-1 targets were distributed across each sector of the repetitive temporal pattern and were not statistically enriched for any phase of expression. The distributed associations of these TFs and their varied targets combined with the constitutive expression of BLMP-1 and ELT-3 throughout each larval stage suggest that BLMP-1 and ELT-3 are associated with their targets throughout the transcriptional cycle regardless of transcriptional class of the target gene or their timing of expression.

### Conserved sequences of the upstream regulatory regions of *lin-4* are necessary and sufficient for high amplitude expression in hypodermal tissues

Many of the heterochronic phenotypes associated with a null mutant of *lin-4, lin-4(e912)*, can be partially rescued by a high-copy transformation of a 693bp genomic fragment that contains approximately 500bp of upstream putative regulatory sequence (Lee et al., 1993). Analysis of the RNAs that are produced from the *lin-4* locus indicate transcription of *pri-lin-4* is initiated at atleast five TSS upstream of the *lin-4* gene (Bracht et al., 2010). Of these TSSs, three occur upstream of the minimal rescuing fragment, suggesting that additional regulatory information may be required to control normal aspects of *lin-4* transcription. These upstream regions are also associated with the active histone modifications H3K4me3 and open chromatin as measured by ATAC-seq assays (Figure 4A) (Daugherty et al., 2017; Janes et al., 2018). We compared the syntenic sequences of five additional nematode species to determine if regions of the F59G1.4 ninth intron were conserved through evolutionary selection. This analysis revealed at least three discrete regions upstream of each *lin-4* gene that were disproportionately conserved when compared to similarly sized intronic and intragenic regions (Conserved Elements (CE) A, C and D). Examination of our ChIP-seq data indicate that each of these regions is enriched for BLMP-1 binding (Figure 4A, Table S1). We did not detect a strong enrichment of ELT-3 bound to the putative *lin-4* regulatory sequences though multiple predicted ELT-3 consensus binding site sequences are found in this region (Figure 4A).

**Figure 4.**
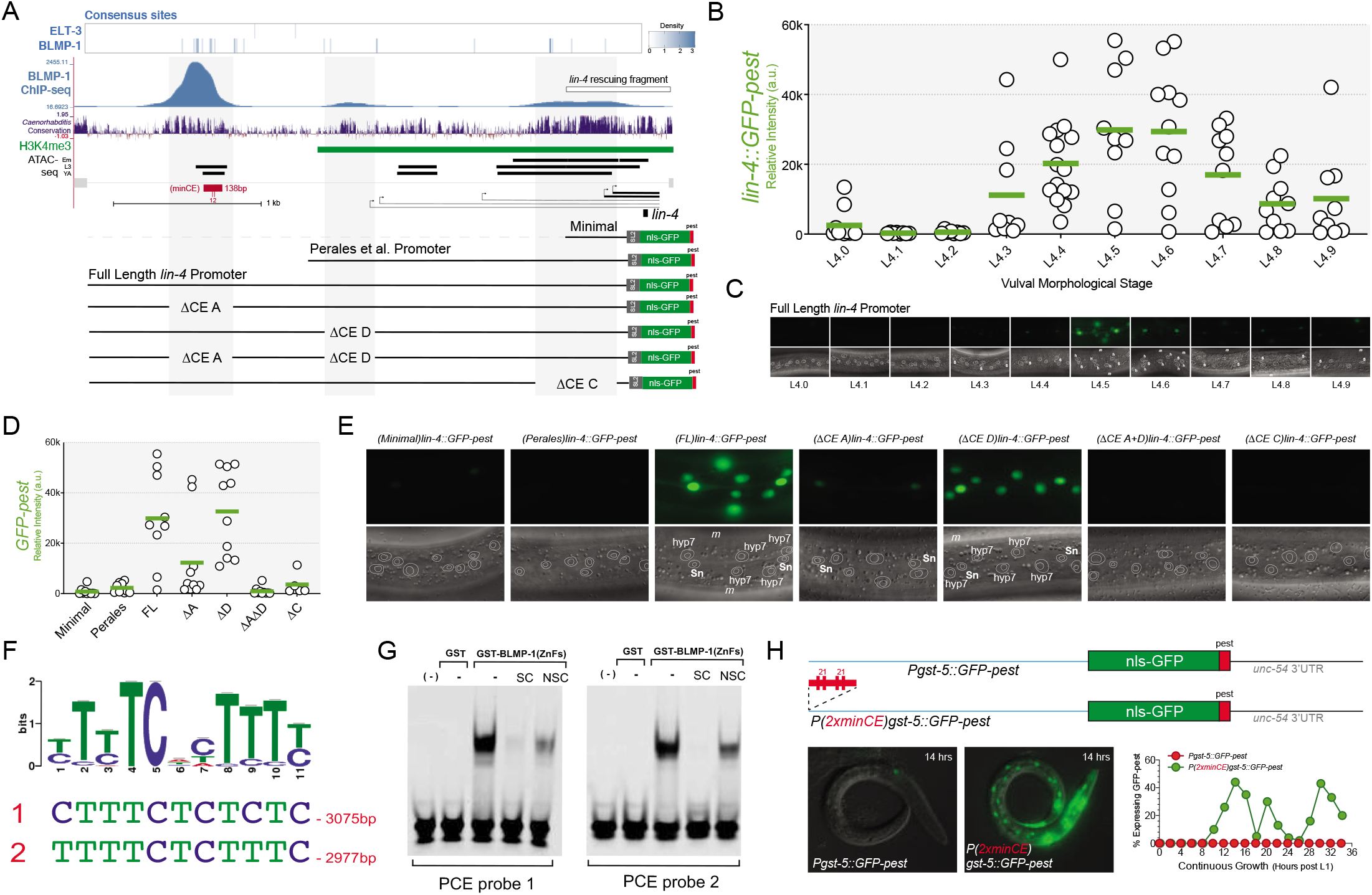
Conserved elements in the *lin-4* promoter are both necessary and sufficient for high-amplitude expression in hypodermal tissues. **(A)** A browser track overlaying the predicted BLMP-1 consensus binding sites, the BLMP-1::GFP ChIP-seq signal from L1-staged larvae, and conservation of these sequence features found in *C. elegans, C. brenneri, C. tropicalis, C. remanei, C. briggsae* and *C. japonica* near the *lin-4* locus and promoters used in the expression analysis. Green bars indicate H3K4me3 ChIP-seq data tracks (indicative of active promoters near transcriptional start sites). Black bars indicate the localization of open chromatin as measured from ATAC-seq experiments (Daugherty et al., 2017). **(B)** Representative measurements of full-length *lin-4::GFP-pest* expression levels in lateral seam cells. The fluorescent intensity of five seam cells per animal were averaged and 8-15 animals were measured per indicated L4 morphological substage. **(C)** Representative fluorescence and DIC images of the full-length *lin-4::GFP-pest* transcriptional reporter in the hypodermal cells of developing L4-staged animals. L4-stage numbers correspond to the defined sub-stages of vulval morphological development (Sn = seam cell nuclei, h = hyp7 cell nuclei, and m = muscle cell nuclei) (Mok et al., 2015). **(D)** Measurements of seam cell expression levels for the various *lin-4::GFP-pest* reporter constructs outlined in panel A at the L4.5 stage of development. (n= 5-10 animals per transgenic strain) **(E)** Representative images of each transcriptional reporter outlined in D. Individual labels defining cell types are the same as in panel C. **(F)** Position Weight Matrix of the consensus BLMP-1 binding site calculated from ChIP-seq data outlined in Figure 3A and two sequences located in the minCE element that conform to this consensus. **(G)** Gel shift experiments demonstrating that a GST fusion protein harboring the 5 ZnF domains of BLMP-1 (but not GST alone) can bind to the two sequences outlined in panel F. **(H)** Comparison of the GFP-pest expression patterns of *gst-5::GFP-pest* or *(2xminCE)gst-5::GFP-pest* in late L1-stage animals. The pulsatile expression of the *(2xminCE)gst-5::GFP-pest* expression correlates with a single pulse in the middle of each larval stage. For the developmental time course, a minimum of forty animals expressing *gst-5::GFP-pest* or *(2xminCE)gst-5::GFP-pest* were scored.

To test the functional relevance of these elements, we constructed a transcriptional GFP-pest reporter that harbors a SL2 splice acceptor sequence directly upstream of the GFP open reading frame (Figure 4A; Full Length (FL)). This feature enables each reporter to generate a single, defined mRNA transcript from each of the various candidate TSSs. This reporter is robustly expressed in a variety of cell types including the hypodermis, pharynx, body wall muscle, and ventral neurons on the surface of larva. Expression of the *lin-4::GFP-pest* reporter occurs once per larval stage (similar to other characterized transcriptional reporters) in these cell types indicating that this sequence contains the information to drive temporally and spatially restricted expression in multiple cell lineages. To characterize the temporal expression patterns in detail, we measured the levels of *lin-4::GFP-pest* expression in lateral seam cells at each of the morphologically defined stages of L4 development (Mok et al., 2015). These experiments revealed that *GFP-pest* expression is highly pulsatile with expression peaking at the L4.5 and L4.6 stages (Figure 4B and C). Expression in other cell types (neuronal, pharynx, muscle and hyp7 cells) also peak during these larval substages with different expression trajectories (Data not shown). Specifically, muscle cells exhibit a broader (longer) transcriptional pulse compared to those observed in seam cells. Expression in hyp7 and neuronal cell types exhibit a shorter pulse of transcription either muscle or seam cells (Figure 4B).

We then compared the expression patterns we observed using the full-length construct to previously published reporters or variants of the full-length reporter that lack individual or combinations of the conserved sequence elements outlined above. Each of the reporters express pulses of GFP-pest expression in similar cell types as the full-length version (Bracht et al., 2010; Perales et al., 2014). However, peak expression in the hypodermal cells was typically 5-20-fold higher from the full-length reporter compared to the other derivative reporter genes (Figure 4D). Deletion analyses of the full-length construct indicate that CE A, harboring sequences that bind BLMP-1 *in vivo*, are primarily responsible for this increased transcriptional output as reporters that lack this element exhibit residual expression to a similar level as the minimal rescuing fragment (Figure 4D and E).

Using the ChIP-seq data sets for BLMP-1::GFP, we identified a consensus sequence for BLMP-1 that is enriched in the CE A regulatory sequence (Figure 4A and C). A recombinant GST translational fusion of the five BLMP-1 zinc fingers (ZnFs), GST-BLMP-1(ZnFs), but not GST alone bound several sequences found in CE A element that conform to this consensus. This binding was diminished by the addition of cold competitor DNA harboring consensus BLMP-1 binding sites but not by a similar sized, non-specific target DNA (Figure 4G). To determine if these elements function as a transcriptional enhancer, we cloned a 138bp fragment from the CE A element, called minimal CE (minCE) (Figure 4A and H), into the 5’ regulatory sequences of the *gst-5* gene encoding the *C. elegans* ortholog of human HPGDS (hematopoietic prostaglandin D synthetase). RNA-seq experiments demonstrate that *gst-5* mRNAs are expressed throughout larval development in a variety of tissues (including the hypodermis) and exhibit a non-cyclical or flat expression pattern (Cao et al., 2017; Hendriks et al., 2014; Kim et al., 2013). A transcriptional reporter bearing 2.9kb of the *gst-5* upstream regulatory sequences drives a low level GFP-pest expression in a number of larval cell types including the pharynx, intestine, neurons and hypodermis and this expression pattern remained below the threshold of imaging during normal development (Figure 4H). In contrast, the transcriptional reporter that included two copies of the minCE in the reverse orientation (reversed compared to the orientation within the *lin-4* enhancer element) drove high amplitude, periodic expression of GFP-pest exclusively in hypodermal cells (Figure 4H). Expression of the *(2xminCE)gst-5::GFP-pest* reporter in *blmp-1(0)* mutant animals resembled the expression of the simple, *gst-5::GFP-pest* reporter in wild-type animals (Figure S4), consistent with the hypothesis that this minimal sequence from the *lin-4* upstream region functions as both a spatial and temporal enhancer.

### BLMP-1 controls the transcriptional output of its targets

A large fraction of BLMP-1 and ELT-3 targets are expressed in a cyclical fashion (approximately 26% and 23%, respectively) (Figure 3D and Table S2). We reasoned that targets that exhibit this highly dynamic form of gene expression could provide an opportunity to determine the effects of BLMP-1 and/or ELT-3 binding on transcriptional dynamics in more detail. To quantify changes in expression dynamics, we obtained or constructed integrated transcriptional GFP-pest reporters that fit two criteria. First, we identified candidate genes that are predicted to be targets of BLMP-1 and/or ELT-3 as measured by ChIP-seq (Figure 3). Second, we selected genes that are predicted (by RNA-seq experiments) to be dynamically expressed in hypodermal tissues (Cao et al., 2017). We therefore constructed or obtained multi-copy integrated transcriptional GFP-pest reporters for *(2xminCE)gst-5, moe-3, mlt-10*, and *ZK180.5* and a single copy version of a transcriptional GFP-pest reporter for *C02E7.6* integrated on chromosome II. To characterize the temporal expression patterns of these reporters, we assembled a series of measurements that spanned the morphologically defined stages of L4 development. As shown in Figures 5A-C, each reporter is expressed in a pulsatile expression pattern with *(2xminCE)gst-5* and *C02E7.6 reporters* exhibiting a peak of expression in the mid-L4 stage, and the *moe-3* transcriptional reporter reaching maximal amplitude near the end. We next aimed to determine how mutations in *blmp-1* and/or *elt-3* altered these expression patterns. GFP-pest expression levels in seam cells were then quantified in each genetic context at a developmental stage where reporter expression peaks in wild-type animals. As illustrated in Figure 5A and B, *blmp-1(0)* mutations dampen the expression of both *(2xminCE)gst-5* and *C02E7.6* reporters by approximately 1.7-fold. In contrast, *elt-3(0)* mutations had little effect on the transcriptional output of either of these two reporters. Furthermore, the decrease in seam cell expression for the *(2xminCE)gst-5* and C02E7.6 reporters was not further decreased in *blmp-1(0); elt-3(0)* double mutants, consistent with a lack of ELT-3 binding sites in these promoters. In contrast, examination of *moe-3::GFP-pest* expression in various mutant backgrounds indicate that activity of both *blmp-1* and *elt-3* are required to maintain high transcriptional output for this gene in the hypodermis (Figure 5C).

**Figure 5.**
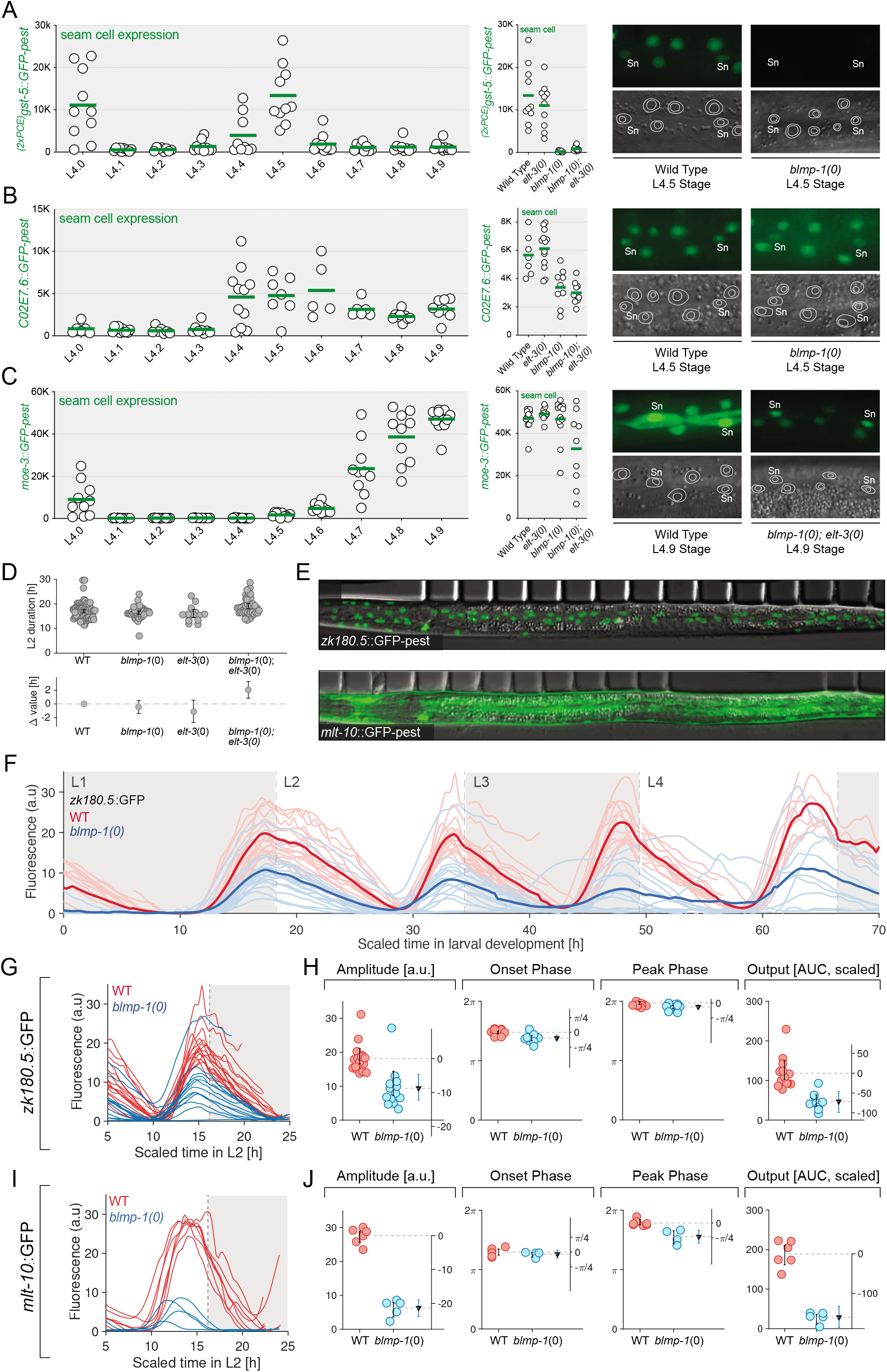
*blmp-1* modulates the amplitude and transcriptional duration of their cyclically expressed targets. **(A-C)** Quantification of GFP-pest expression from the *(2xminCE)gst-5, C02E7.6* and *moe-3* transcriptional reporters in L4-staged animals. Top panel for each reporter indicates the average fluorescent intensity of GFP-pest expression in the lateral seam cells for each morphologically defined L4 stage (*(2xminCE)gst-5* = L4.5, C02E7.6 = L4.5, and *moe-3* = L4.9) (n = 7-14 animals)(Sn = seam cell nuclei). **(D)** Estimation plot (Ho et al., 2019) showing the duration of second larval stage (L2) for WT (N=92), *blmp-1(0)* (N=46), *elt-3(0)* (N=15) and *blmp-1(0); elt-3(0)* animals (N=50). Upper row: Grey circles indicate L2 duration, measured by observing cuticle shedding (molts) during long-term live imaging using a microfluidics device (Keil et al., 2016). (see Methods). Black error bars indicate mean and standard deviation. Lower row: Mean bootstrapped difference values compared to WT and 95% confidence intervals of bootstrap distributions (black error bars). **(E)** Merged DIC and fluorescence micrographs (after worm axis straightening (Keil et al., 2016)) showing reporter expression at peak intensity in the L2 stage. **(F)** *ZK180.5::GFP-pest* reporter intensity, averaged over the entire animal throughout development as a function of time for WT (light red, N=23, dark red average) and *blmp-1(0)* mutant (light blue, N=13, dark blue average). Time courses are scaled such that molts (dashed grey lines) of individual animals align with average WT molting timings. **(G)** Inset of (F), showing the fluorescence intensity during the L2 stage of development. **(H)** L2 peak fluorescence intensity (left most), onset phase of reporter expression (left middle), peak phase of reporter expression (right middle) and Output (AOC = area under the curve) (right most) for the *ZK180.5::GFP-pest* reporter in WT (red circles, N=21) and *blmp-1(0)* mutant (blue circles, N=13). Black triangles indicate mean bootstrapped difference values, blue error bars indicate 95% confidence intervals of bootstrapped difference distribution. **(I,J)** As (F, G) but for *mlt-10::GFP-pest* (WT, N=7; *blmp-1(0)*, N=5).

We next sought to correlate dynamic changes in gene expression to other aspects of larval development in more detail. To accomplish this, we performed long-term, live imaging of developing larvae in a microfluidic device (Keil et al., 2016). This imaging strategy enabled us to continuously monitor gene expression in individual animals and also correlate these changes with other cellular and developmental milestones. First, we compared overall developmental pace between WT, *blmp-1(0), elt-3(0)* and *blmp-1(0);elt-3(0)* animals in the microfluidic chambers by assessing the time between the successive ecdysis (cuticle shedding). While both WT and mutant animals displayed variability of larval stage duration within each genotype, neither *blmp-1(0)* nor *elt-3(0)* mutations (or combination of these mutations) affected the average duration to a substantial degree (Figure 5D and Figure S5). However, our live imaging experiments revealed a partially penetrant larval lethality phenotype, in particular in the L3 stage, that was not apparent on standard, solid NGM plates.

The fact that the larval stage durations are largely preserved between WT and *blmp-1(0)/elt-3(0)* mutants allowed us to directly compare average quantitative temporal features of transgene gene expression dynamics in each genetic background. To this end, we directly monitored patterns of two *GFP-pest* reporters, *ZK180.5::GFP-pest* and *mlt-10::GFP-pest*, that are exclusively expressed in the hypodermis (Figure 5E) and whose putative regulatory elements are bound by BLMP-1 and ELT-3 (Figure 3 and Figure S6). To compare the expression patterns between animals, we extracted individual animal fluorescence time course measurements and rescaled them such as to align the timing of molting events of each animal (see Methods) to the average WT molting times (Figure 5F). This procedure enabled us to directly compare specific features between each expression trajectory including amplitude, transcriptional onset (the time within each transcriptional cycle in which transcription begins), peak phase (the point within each cycle in which the peak expression is reached), and the cumulative accumulation of GFP-pest expression (output). Consistent with other *blmp-1*- or *blmp-1; elt-3*-dependent reporters, cyclical transcription of *ZK180.5::GFP-pest* and *mlt-10::GFP-pest* was maintained throughout larval development indicating that *blmp-1* (or *elt-3*) are not essential for generating pulsatile patterns of transcription (Figure 5 F-J and Figure S6). The primary difference between the expression trajectories of wild type and mutant animals was the drastic reduction in the amplitude of individual traces in *blmp-1(0)* mutants. Surprisingly, the reduction of transcriptional output was not accompanied by any substantial change in the timing of transcriptional induction (Figure 5H and J and Figure S6). Ultimately, the reduction of transcriptional output (as measured by total cumulative GFP-pest expression during each pulse) correlates with a reduction in the relative duration of expression in mutant animals (Figure 5H and J and Figure S6). These results indicate that BLMP-1 functions to amplify the baseline pulsatile transcriptional dynamics that are likely generated by other gene regulatory mechanisms.

### The *lin-4* locus adopts a BLMP-1-dependent, active chromatin structure in hypodermal cells

Previous studies indicate that BLMP-1 physically and functionally interacts with multiple chromatin remodeling factors (e.g., SWI/SNF subunits) and that BLMP-1 binding sites are enriched in open chromatin regions (Fong et al., 2020; Daugherty et al., 2017; Janes et al., 2018). We hypothesized that BLMP-1 may function to potentiate transcription through chromatin remodeling near its binding sites which would potentiate the binding of additional TFs. This model would account for the dependency of the diverse transcriptional targets of BLMP-1 that differ in their temporal pattern of expression (phase) during the repetitive transcriptional cycle and would be mechanistically similar to the activities of other TFs (including pioneer factors) that remodel chromatin accessibility prior to the onset of major transcriptional induction in *C. elegans* (Cochella and Hobert, 2012; Hsu et al., 2015; Patel and Hobert, 2017).

We therefore sought to measure how BLMP-1 activity may alter chromatin dynamics at a specific locus with spatiotemporal resolution. We utilized an established chromosome-tagging strategy that has been employed to visualize both the localization of transgenic arrays and dynamics of chromatin compaction in living animals (Cochella and Hobert, 2012; Fakhouri et al., 2010; Meister et al., 2010). Transgenes containing the *lin-4(FL)::mCherry-pest* ORF flanked by tandem copies of the bacterial lac operator (lacO) sequences were integrated to a single chromosomal site and then crossed into a strain harboring a soluble, ubiquitously expressed GFP-tagged lac repressor (LacI) fusion that recognizes lacO sequences with high affinity (Patel and Hobert, 2017). The enabled us to monitor the *lin-4* locus in live animals while simultaneously monitoring transcriptional output of the *lin-4* enhancer/promoter (Figure 6A). Animals harboring these transgenes exhibit two foci in each somatic nucleus indicating the stable, single integration site of the *lin-4::mCherry/LacO* transgene. The intranuclear GFP-LacI foci found in freshly hatched L1-staged animals can be grossly categorized into two basic classes: Class 1) a punctate or “ball-like” pair of GFP-LacI foci that are found in a majority of somatic cells and Class 2) pairs of “puffed” foci that are found in the nuclei of hypodermal cells (Figure 6B). This lineage-specific decompaction of the *lin-4* locus is surprising for multiple reasons. First, *lin-4* loci appear to be decompacted only in hypodermal tissues where BLMP-1 is normally expressed. Second, chromatin decompaction is typically associated with active transcription in a variety of systems (Dietzel et al., 2004; Tumbar et al., 1999; Yuzyuk et al., 2009) and the decompaction of the *lin-4* loci in freshly hatched L1-stage animals precedes the transcriptional induction of *lin-4* miRNAs by approximately 12 hours (note a lack of *mCherry* expression in Figure 6B).

**Figure 6.**
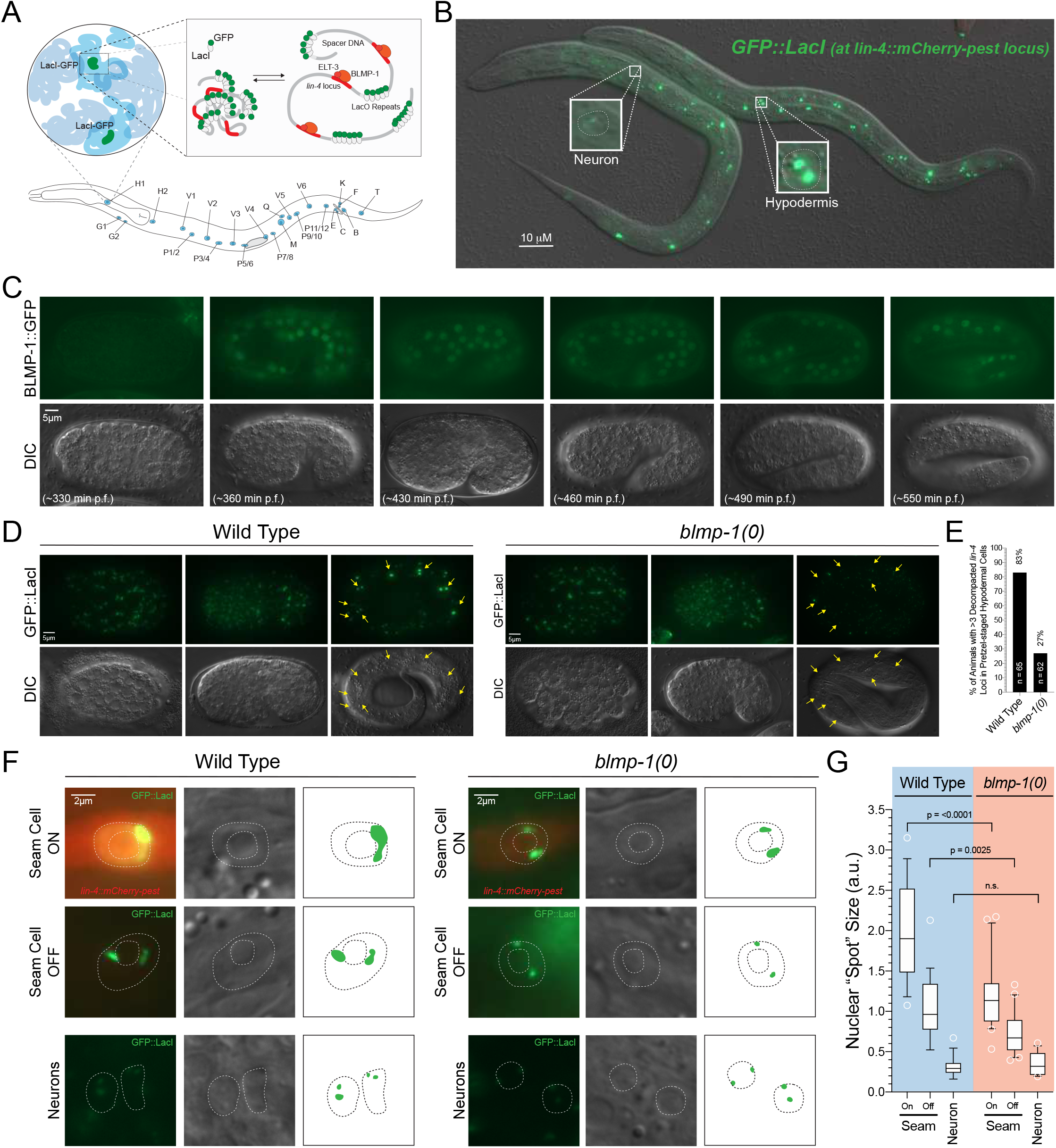
The *lin-4* locus is maintained in an open chromatin state in hypodermal cells in a *blmp-1*-dependent manner. **(A)** Schematic of the transgenic array used for visualization of the *lin-4* locus in living animals. **(B)** Representative image of L1-staged animals expressing the GFP-lacO-labeled *lin-4* locus. **(C)** Analysis of the expression pattern of an endogenously GFP-tagged translational fusion of BLMP-1 indicating that expression begins in hypodermal cells around the bean stages and remains stably expressed throughout embryonic development. **(D)** Pictographs depicting the compression level of the *lin-4* locus/GFP-LacI foci in differentially staged wild-type and *blmp-1(0)* mutant embryos. Yellow arrows indicate the location of hypodermal nuclei. **(E)** Quantification of the percentage of wild-type or *blmp-1(0)* late-stage (pretzel) embryos that exhibit decompacted *lin-4* loci in hypodermal cells. **(F)** Close-up images of representative seam cell and neuronal nuclei from L3-staged wild-type and *blmp-1(0)* mutant larva. GFP images are the maximum intensity projections obtained for each relevant nuclei. **(G)** Quantification of the sizes of the nuclear GFP-LacI foci in various cell types of wild-type and *blmp-1(0)* mutant animals. Boxes and median line indicate the interquartile range. Whiskers cover 10-90th percentile. Circles represent outliers. Brackets indicate statistically significant differences in “puffs size” calculated from a two-tailed chi-square analysis.

To test whether BLMP-1 expression correlates with target loci decompaction, we performed two types of analyses. First, we characterized when BLMP-1 expression begins in embryos. BLMP-1::GFP can be detected as early as the bean stage (approximately 365 minutes post fertilization) in most developing hypodermal cells (Figure 6C). This tissue-specific expression pattern remains throughout the rest of embryonic development (Figure 6C). We next aimed to determine how the timing of BLMP-1 expression correlates with the decompaction level of the *lin-4* loci by examining changes in GFP-LacI foci. From the onset of GFP-LacI expression to the early gastrulation stage of embryogenesis, most somatic *lin-4* GFP-LacI foci are decompacted, consistent with a general lack of condensed chromatin and reduced histone H3 lysine 9 (H3K9) methyltransferase activity in early embryos (Mutlu et al., 2019; Mutlu et al., 2018; Towbin et al., 2012; Yuzyuk et al., 2009). By the end of gastrulation (approximately 330 minutes post fertilization) the foci in most somatic cells are equally condensed consistent with the onset of large-scale heterochromatin formation in somatic cells and a reduction in transcriptional plasticity (Mutlu et al., 2019; Mutlu et al., 2018; Yuzyuk et al., 2009). At roughly the same time as the onset of BLMP-1 expression in embryonic hypodermal cells, the GFP-LacI foci begin to decompact in posterior seam cells (Figure 6D). The cell type-specific decompaction of *lin-4* loci in hypodermal tissues is complete for a majority of wild-type animals by the end of embryogenesis while the same foci in other somatic cell types remain tightly compressed (Figure 6D). GFP-LacI foci dynamics were similar in wild-type and *blmp-1(0)* animals during early embryogenesis. In contrast to the late embryonic decompaction of *lin-4* loci in wild-type animals, hypodermal GFP-LacI foci (and foci in other somatic nuclei) in *blmp-1(0)* mutants remained condensed (Figure 6D and E). We conclude that BLMP-1 is required for decompaction of the *lin-4* loci in hypodermal cells during embryogenesis. Furthermore, this decompaction occurs at a point in development where *lin-4* transcription is being actively repressed by multiple FLYWCH proteins (Ow et al. 2008).

We next aimed to directly monitor any dynamic changes in “puffs” sizes within the transcriptional cycle of *lin-4* during larval development and determine if these changes are dependent on BLMP-1 activity. In head ganglion cells, derived from a cell lineage that never express the *lin-4pro::mCherry-pest* transgene, the two *lin-4pro::mCherry-pest* loci are constitutively compacted (Figure 6F and G). To measure changes in cells that do express *lin-4* miRNAs, we focused our analysis on developing lateral seam cells for multiple reasons. First, seam cells constitutively express BLMP-1 and dynamically express the *lin-4pro::mCherry-pest* reporter at specific phases of each larval stage (Figure 1B and Figure 4A). Second, in contrast to hyp7 syncytial cells that endoreduplicate their nuclear DNA content at each division resulting in a stable ploidy of 4n, seam cells replicate their nuclear DNA once per division at specific, defined phases of the larval molt cycle (Hedgecock and White, 1985). To ensure accuracy in both the timing of *lin-4::mCherry-pest* expression and seam cell divisions, we specifically measured the compaction status of the *lin-4::mCherry-pest* loci in late L3-staged animals where morphological features of the vulval cell lineage, gonad morphology, and *mCherry-pest* expression could be used to compare animals of both genotypes at similar developmental stages. We found that the GFP-LacI “puffs” size in non-transcriptionally active lateral seam cells was typically three times larger than the those found in neurons (Figure 6F and G). Later, during the L3 lethargus, when *lin-4pro::mCherry-pest* is expressed, we observed that the average size of the GFP-LacI “puffs” further increase in size (approximately an additional 2-fold when compared to non-transcriptionally active seam cells) (Figure 6F and G). This suggests that the *lin-4* locus is constitutively decompacted (compared to the same locus in neuronal cells) and the process of transcribing this locus further relaxes chromatin architecture. We then asked if chromatin decompaction at these loci was dependent on BLMP-1 activity by examining the same features in *blmp-1(0)* animals. A comparison of the GFP-LacI “puff” sizes in head neurons indicates that they are equally compressed in wild-type and *blmp-1(0)* animals (Figure 6F and G). In contrast, GFP-LacI foci, in both non-transcribing and *lin-4* expressing cells, are significantly reduced in size compared to those observed in stage-matched wild-type animals (Figure 6F and G). These experiments demonstrate that the chromatin structure of the *lin-4* locus is dynamically remodeled in seam cells and that BLMP-1 is required for decompaction of the *lin-4* locus in hypodermal cells prior to transcription in wild-type, larval animals.

### BLMP-1 and ELT-3 are required to resume normal temporal patterning after nutrient deprivation

Development of *C. elegans* larvae in laboratory conditions is both rapid and continuous due to an abundant, controlled food supply and standardized growth conditions. In contrast, the natural environment in which *C. elegans* larvae normally live is far more diverse and dynamic leading to a more punctuated developmental trajectory (Frezal and Felix, 2015). Acute food removal induces a defined developmental diapause that can occur immediately after each larval molt (Schindler et al. 2014). The diapause is part of a developmental checkpoint that is distinct from dauer development and controlled by insulin signaling (Schindler et al. 2014). When the insulin signaling rapidly inactivated (i.e. during acute starvation), animals finish the current larval stage, arrest all somatic cell movements and divisions, and halt the expression of molting-specific genes (Schindler et al., 2014).

To determine if global aspects of cyclical transcription are altered during acute starvation, we monitored the expression of four transcriptional reporters that exhibit distinct phases of expression (Figure 5). These reporters include targets of BLMP-1 and ELT-3 (*(minCE)gst-5::GFP-pest, ZK180.5::GFP-pest*, and *mlt-10::GFP-pest*) as well as a transcriptional reporter of *blmp-1, since* ChIP-seq data indicates that it is expressed in a cyclical expression pattern (Hendriks et al., 2014; Kim et al., 2013) and BLMP-1 activity regulates its own expression in an autocatalytic fashion (Figure S7). We grew populations of L1-staged animals under nutrient-rich conditions for 22 hours and then one half of each sample was allowed to continue development in nutrient replete conditions while the other half was acutely starved (Figure 7A). Acute food removal results in greater than 90% of transgenic animals arresting development with undivided vulval precursor cells after 24 hours indicating that larval development has been paused (Schindler et al., 2014). During the starvation period, we monitored GFP-pest expression in fed and starved populations. In contrast to the cyclical expression of each reporter under nutrient replete conditions, GFP-pest expression for each of the reporters was extinguished during starvation conditions with similar kinetics (Figure 7B). This transcriptionally inactive state was maintained throughout the starvation period. Re-fed animals resumed normal, cyclical expression patterns within a few hours after food exposure (Figure 7B). These experiments demonstrate that oscillatory transcription is inhibited by the starvation-induced developmental diapause and that wild-type animals can re-initiate these transcriptional patterns in a nutrient-dependent manner.

**Figure 7.**
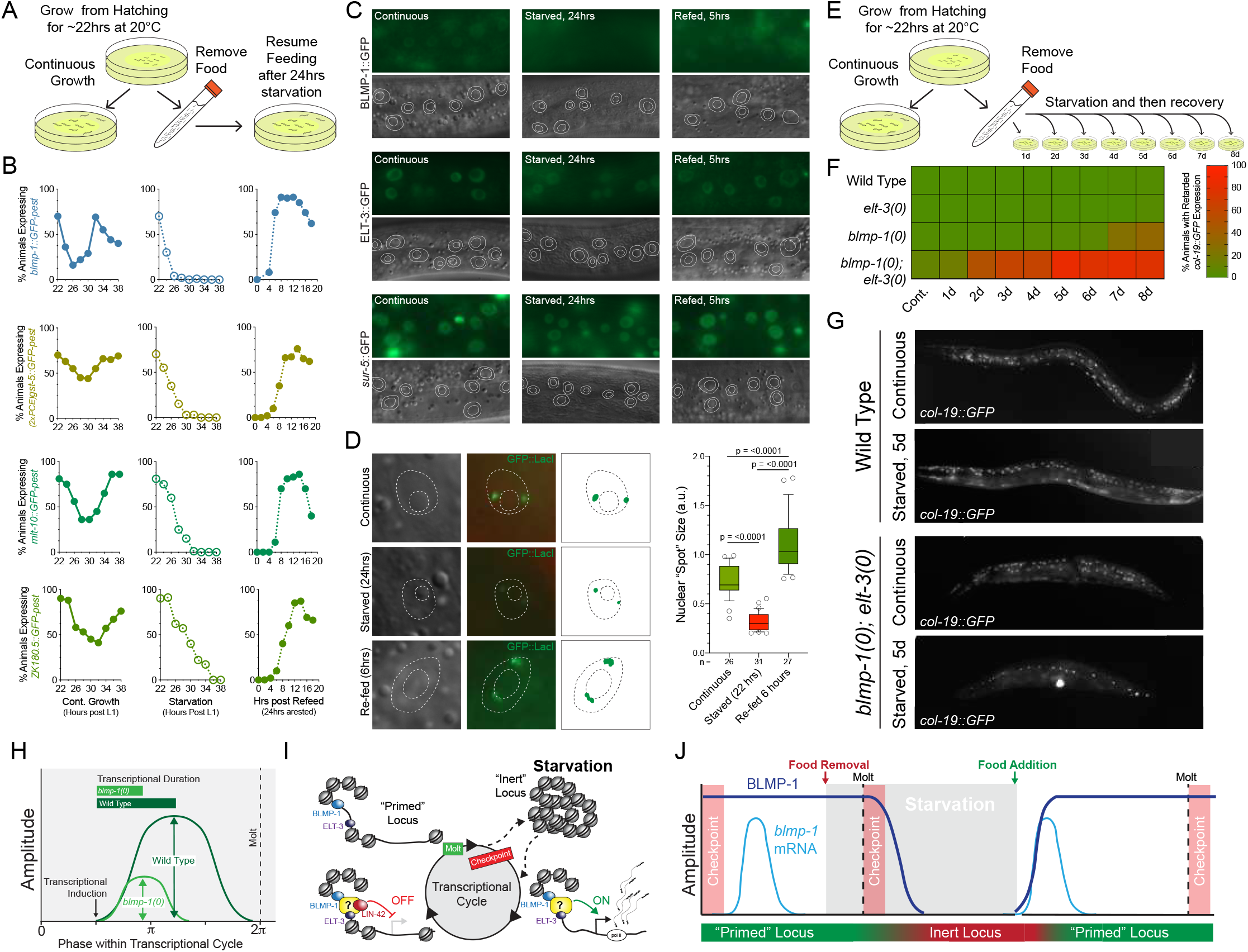
Dynamic gene expression is halted during nutrition-mediated developmental arrest and BLMP-1 and ELT-3 are essential for the recovery of normal temporal patterning after starvation. **(A)** A schematic of the starvation experiment used to measure nutrient-depended changes in the expression patterns of cyclically expressed genes. **(B)** Continuous cycling of gene expression is arrested during starvation conditions and is reinitiated in a coordinated manner when animals resume development. For each timepoint, 70-200 animals were scored. **(C)** BLMP-1::GFP expression is down-regulated during starvation conditions and expression is rapidly resumed when animals reinitiate development. **(D)** The *lin-4* locus is compacted during starvation conditions in lateral seam cells. **(E)** A schematic of the starvation-mediated arrest and re-feeding experiment used to measure the ability of animals to resume normal temporal patterning after starvation. **(F)** The resumption of normal temporal patterning after starvation requires *blmp-1* and *elt-3* activity. Defects in temporal development were monitored by measuring the *col-19::GFP* expression phenotypes in re-fed animals after they resumed growth for 24hours after the indicated period of starvation. (n = >100 per condition) **(G)** Representative pictographs of the *col-19::GFP* phenotypes in wild-type and *blmp-1(0); elt-3(0)* mutants after animals have recovered from 5 days of starvation. **(H)** A diagram outlining the expression features of a single transcriptional cycle in wild-type and *blmp-1(0)* animals. **(I)** A model for how BLMP-1 and ELT-3 function to regulate transcriptional output during larval development. **(J)** A diagram outlining the changes of *blmp-1* mRNA and BLMP-1 protein levels during normal grown and during starvation conditions.

BLMP-1 expression is actively regulated post-translationally by a conserved F-box protein, DRE-1, that mediates the ubiquitination of target proteins which are degraded by the proteasome (Horn et al., 2014). Given that *blmp-1* is cyclically transcribed and its transcription is inhibited in nutrient depleted conditions (Figure 7B), we hypothesized that the constitutive expression of BLMP-1 during normal development may be altered during starvation. To test this, we repeated the nutrition-mediated arrest experiments outlined above in strains harboring BLMP-1::GFP and ELT-3::GFP translational fusions. The expression patterns of each translational fusion were compared to a constitutively expressed monomeric GFP transgene (*sur-5::GFP*) to control for potential changes in GFP half-life under starvation conditions. Surprisingly, BLMP-1::GFP expression was diminished during starvation with >50% of animals loosing detectible BLMP-1::GFP expression after 12 hours of starvation. In contrast to the down-regulation of BLMP-1::GFP, most starved animals retained ELT-3::GFP and soluble GFP expression throughout the starvation period with only a mild reduction after 24 hours (Figure 7C). As with the transcriptional reporter for *blmp-1* expression, BLMP-1::GFP expression was detectible again in most animals within 2 hours or re-feeding and resumed normal, pre-starved expression levels after 5 hours (Figure 7C). Similar changes in *blmp-1* mRNA and BLMP-1 protein expression were seen in freshly hatched and starved L1 (Figure S8). We conclude that BLMP-1 expression, both transcriptionally and post translationally, is regulated by nutrient availability.

To examine if BLMP-1-dependent chromatin remodeling is also modulated by nutrient-dependent signaling, we monitored compaction of *lin-4* locus in animals that have entered the acute starvation-dependent developmental diapause. At the mid-L3 stage (first VPC division), a developmental time point that precedes *lin-4* transcription during the L3 stage the *lin-4* locus was decompacted (Figure 7D). We then rapidly removed the bacteria food source to induce the L4-specific developmental diapause. After 22 hours of starvation, >90% of animals arrested at the invagination stage of vulval development (L4.1-L4.2) which precedes fusion between the anchor cell and vulval precursor cells on the ventral surface of the L4 larva (Mok et al., 2015). Analysis of lateral seam cell GFP-LacI puffs sizes in starved animals demonstrated that the *lin-4* loci in seam cells were dramatically compacted when compared to the same loci in continuously developing animals (Figure 7D). Re-plating of these starved animals lead to the rapid decompaction of the *lin-4* locus at a timepoint with BLMP-1 expression has resumed (Figure 7D). Interestingly, the level of decompaction is greater in recovering animals that those found in non-starved, continuously developing animals (Figure 7D). We also monitored *lin-4::mCherry/LacO* loci in the seam cells of starved, L1-stage animals and also found that loci compacted in these starvation conditions (Figure S8). In sum, these experiments indicate that dynamic chromatin rearrangements are modulated by nutrient sensing.

Changes in BLMP-1 expression and the dramatic compaction and decompaction of BLMP-1 target chromatin during starvation and recovery, respectively, suggests that BLMP-1 could play an essential role in modulating transcriptional output of critical developmental genes in dynamic environments. To determine whether animals that lack *blmp-1* and/or *elt-3* can robustly adapt to starvation, we repeated the food experiments as described in Figure 7A but maintained the animals in starvation conditions for increasing periods of time. At the end of the starvation period, animals were re-plated on normal food and monitored for developmental phenotypes by scoring changes in *col-19::GFP* expression when the animals had reached adulthood (Figure 7E). In these experiments, wild-type and *elt-3(0)* larvae re-initiate normal temporal patterning after prolonged bouts of starvation (Figure 7E). In contrast, a minor fraction of *blmp-1(0)* mutants failed to express *col-19::GFP* in hyp7 cells, a phenotype associated with weak reiterative heterochronic mutants (Abbott et al., 2005; Hammell et al., 2009). The relatively mild *blmp-1(0)* phenotypes were dramatically enhanced in *blmp-1(0); elt-3(0)* double mutants. These results indicate that BLMP-1 and ELT-3 are essential for resuming normal temporal patterning after continuous development has been interrupted by starvation.

## Discussion

### Features of the oscillatory expression program impose constraints on modulating transcriptional output

Oscillatory patterns of gene expression are a ubiquitous feature of developmental systems where they orchestrate repetitive biological functions and/or anticipate recurring environmental conditions (Hasty et al., 2010). While most GRNs that program cyclical patterns of gene expression utilize a basic negative feedback structure, specific features of their architecture define the precision of the timekeeping mechanism and the ability to modulate and/or preserve features of dynamic transcription. For instance, the periodicity of circadian gene expression persists in the absence of external cues over a wide range of temperatures, yet changes in lighting or nutrition can alter the phase of gene expression or the amplitude of clock target gene transcription, respectively (Shigeyoshi et al., 1997; Trott and Menet, 2018). The genetic control of plant root formation also employs a complex feedback loop that generates cyclical transcriptional cycles that induce the formation of lateral roots at periodic positions along the radial surfaces of the developing primary root. The transcriptional output of this GRN is modulated by a hormone, auxin, that amplifies oscillatory transcription in a dose-dependent manner (Moreno-Risueno et al., 2010). Exposure to light is required for maintaining the oscillatory transcriptional patterns at the pre-branch sites (Kircher and Schopfer, 2016). Water availability can also modulate the ability to the root clock to induce lateral root founder cell development. Uneven water distribution on the surface of the developing primary root leads to a suppression of LR development on the dryer root surface (Bao et al., 2014). Importantly, changes in water distribution do not alter the cycling of core root clock components indicating the environment can control clock output at the phenotypic level (Bao et al., 2014).

While the molecular components of the *C. elegans* expression oscillator are not fully known, a detailed characterization of the systems-level behavior of this GRN indicates that two features are hard-wired. First, the biological oscillator is directly tied to the molting cycle and gated by genetically-regulated developmental checkpoints (Hendriks et al., 2014; Schindler et al., 2014). Under replete nutritional conditions, transcriptional cycling is continuous (Hendriks et al., 2014). Removal of food either at hatching or just prior to the termination of the molt results in a regulated arrest of the developmental clock at a precise point within the transcriptional cycle (Figure 7B)(Hendriks et al., 2014). Re-initiation of the transcriptional clock (either from starved L1-stage animals, dauer arrest, or the nutrition-mediated arrest points at each larval stage) occurs at this same point (Hendriks et al. 2014). These features indicate that once initiated, the transcriptional cycle is modular in nature and progresses until the next checkpoint. Secondly, the periodicity of thousands of cyclically expressed transcripts scales inversely with temperature over a wide dynamic range and the relative timing of expression for individual genes within the transcriptional cycle is phase-locked (Hendriks et al., 2014; Kim et al., 2013).

Because the accordion-like, modular structure of this GRN constrains the relative timing of transcriptional events within the cycle, other features of cyclically transcribed genes are likely modulated independently of the clock. This adaption of transcriptional output to various environments would be especially important for genes that are expressed at determinant phases within the transcriptional cycle and function in dosage-sensitive manners. This feature is exemplified by the heterochronic miRNAs that exhibit pulsatile transcriptional patterns at each larval stage and also function to control discrete transitions of temporal cell fate at specific developmental milestones (molting) (Ambros, 2011). It had been previously shown that LIN-42, the *C. elegans* Period ortholog, functions to limit transcriptional output of this class of genes by negatively regulating the overall duration of transcription within each larval stage (McCulloch and Rougvie, 2014; Perales et al., 2014; Van Wynsberghe and Pasquinelli, 2014). In this manuscript, we present evidence that BLMP-1 functions to antagonize LIN-42 activities by increasing the duration of transcription for cyclically expressed genes. As with *lin-42, blmp-1* is not essential for the generation of cyclical expression patterns but plays a modulatory role in controlling the duration and amplitude of transcription (Figure 7G). How the interplay between these two antagonistic modulators of gene expression achieves precise tuning of gene-dosage to confer robustness and adaptivity to post-embryonic developmental programs is an exciting route for further investigation given that BLMP-1 targets the *lin-42* locus (Table S1).

### BLMP-1 couples chromatin de-compaction and the regulation of transcriptional output

Here, we characterize a molecular role for BLMP-1 in regulating the transcriptional output of a number of cyclically expressed genes through a mechanism involving the decompaction of chromatin near its target gene loci. In contrast to the chromatin decompaction that occurs during normal transcriptional activation (Dietzel et al., 2004; Tumbar et al., 1999; Yuzyuk et al., 2009), we demonstrate that BLMP-1-dependent decompaction is temporally separated from the activation of the target gene expression and can occur even when transcription is actively repressed by other mechanisms (e.g. for the *lin-4* gene during embryogenesis (Figure 6B and D)(Ow et al., 2008). We hypothesize that this anticipatory priming mechanism constitutively opens chromatin loci near BLMP-1 binding sites. These “primed” loci would then be accessible to additional TFs whose binding capacity may be normally restricted by nucleosome complexes. Once bound, these factors would induce the phased patterns of gene expression that define the transcriptional cycle for each target gene (Figure 7 G). Because BLMP-1 binding sites are associated with a multitude of genes that are expressed in diverse phases of the oscillatory transcriptional pattern (Figure 3), we speculate that the BLMP-1-dependent “priming” activity is not limited to a single partner TF but likely facilitates the association of several TFs that function at distinct phases of the transcriptional cycle. In the absence of BLMP-1, many of these factors would exhibit a reduced capacity to bind to their cognate binding sites. As a result, *blmp-1(0)* mutants exhibit broad changes in transcriptional dynamics. The diversity of these changes would result in pleiotropic phenotypes whose only common feature would be a change in gene dosage across multiple GRNs. Notably, while our analysis focused on genes whose expression is upregulated in the presence of BLMP-1, the mechanism we propose by which BLMP-1 modulates target transcription is equally capable of inhibiting transcription of target genes (by increasing chromatin accessibility for transcriptional repressors).

The molecular mechanism that we propose for BLMP-1 in temporal gene regulation is analogous to the role that the TF Zelda (ZLD) plays in establishing normal spatial gene regulation during *D. melanogaster* embryogenesis. In this system, ZLD is ubiquitously expressed throughout the embryo where it functions as a pioneer factor to rearrange chromatin near its binding sites (Liang et al., 2008; Foo et al., 2014; McDaniel et al., 2019). Spatially ubiquitous ZLD activity facilitates the transcriptional activation of genes that are co-targeted by the transcription factor Dorsal (DL). Unlike ZLD, DL is expressed in a dorsal-ventral gradient that extends from the dorsal surface of the embryo to the lateral edge. DL functions in a dosage-dependent manner to generate a graded expression of target genes (Rushlow et al. 1989; Roth et al. 1989). Mutation of ZLD binding sites in DL-dependent reporter transgenes results in the dramatic compression of reporter gene expression along the dorsal-ventral axis (Yamada et al, 2019). Importantly, *zld* mutations do not alter DL distribution along the dorsal-ventral axis suggesting that ZLD activity primes DL targets by reorganizing chromatin architecture that would otherwise impair DL binding and function. As with mutations of BLMP-1 in the context of temporal gene expression, *zld* mutations additionally alter the duration of DL target expression (Yamada et al. 2019).

### Regulation of BLMP-1 expression in response to nutrient availability coordinates transcriptional output

We hypothesize that the re-initiation of transcription after starvation elicited by food is essential for animals to re-format the chromatin landscape near their target genes after quiescence and to rapidly modulate the transcriptional output of genes that control temporal patterning. In this manuscript we also show that *blmp-1* expression is regulated by nutrient availability and is important for the coordination of transcriptional output in a variety of conditions. Under nutritionally replete growth conditions, *blmp-1* transcription is both cyclical and auto-regulatory. While *blmp-1* expression at the mRNA level is highly dynamic, BLMP-1 protein expression is highly stable and maintained at relatively constant levels during continuous growth (Hendriks et al., 2014; Horn et al., 2014; Kim et al., 2013). Starvation elicits two main changes in *blmp-1* expression that directly lead to a reduction in the priming activity. First, the cyclical transcription of *blmp-1* (as with most other cyclically expressed genes) is arrested when animals pause development at a starvation-induced developmental checkpoint. Second, BLMP-1 expression is rapidly curtailed during starvation. BLMP-1 expression is likely dampened by the combined reduction in transcription and the constitutive activity of DRE-1 that functions independently of starvation (Horn et al., 2014). This reduction of BLMP-1 expression is coincident with a re-compaction of chromatin near BLMP-1 target genes. The establishment of non-permissive chromatin architecture may be a common feature of quiescent cells as diverse cell types (from yeast to human cells) exhibit these changes. It has also been previously suggested to control inappropriate transcription in the context of development (Evertts et al., 2013; Laporte et al., 2016 Pinon, 1978; Rawlings et al., 2011; Rutledge et al., 2015; Swygert et al., 2019). The regulation of BLMP-1 expression through nutrient availability is also important for the coordination of gene expression after starvation as *blmp-1(0)* and *blmp-1(0); elt-3(0)* animals fail to resume normal temporal patterning after starvation.

Finally, we hypothesize that the active control of BLMP-1 levels/activities may be important for the adaption of global transcriptional patterns to the environment. The separate genetic control of transcriptional timing and transcriptional output would enable gene expression dynamics to be optimized to specific conditions and ensure developmental robustness. Evidence that the levels of BLMP-1 protein are actively regulated are derived from the observation that *dre-1(lf)* phenotypes result in an inappropriate over-expression of BLMP-1 and a coincident increase in precocious developmental phenotypes (Horn et al., 2014). The active control of BLMP-1 protein expression may play an important role in the adaptation of transcriptional output to different temperatures where the periodicity of the transcriptional cycles and developmental pace changes dramatically (Kim et al. 2013; Hendriks et al. 2014). This homeostatic strategy, mediated by controlling the expression of a single pioneer transcription factor, could buffer differences in transcriptional dynamics across the transcriptional cycle by modulating chromatin accessibility.

## EXPERIMENTAL METHODS

See Star Materials and Methods Document.

## ACKNOWLEDGEMENTS

We thank A. Zinovyeva, V. Ambros, L. M. Kutscher, and members of the Hammell laboratory for critical review of this manuscript. We received *C. elegans* strains and recombinant DNAs from S. Gasser, O. Hobert, L. Cochella. The ATIP/Avenir Young Investigator program of the CNRS supported to W.K. W.K. initiated and performed part of this work while being a postdoctoral fellow in the laboratories of Shai Shaham (S.S.) and Eric D. Siggia (E.D.S) at Rockefeller University, supported by NIH grant R35NS105094 to S.S., NSF grant PHY 1502151 to E.D.S. and a postdoctoral fellowship (LT000250/2013-C) from the Human Frontier Science Program (HFSP) to W.K.. Cold Spring Harbor Laboratory, the Rita Allen Foundation, HIH NIGMS R01GM117406 supported C.M.H..

## AUTHOR CONTRIBUTIONS

N.S., W.K., and C.M.H. designed, performed and analyzed most experiments and wrote the manuscript. ChIP-seq experiments were performed by V.E. and S.E.. C.M.H and K.H-M. analyzed sequencing data. Gel shift experiments were carried out by K.D.. Reporter construction and gene expression analysis was carried out by C.M.H., N.S. and K.D.. Microfluidics experiments were carried out by W.K.. Starvation experiments and LacO/LacI experiments were carried out by N.S. and C.M.H..

## SUPPLEMENTAL FIGURES AND TABLE LEGENDS

**Supplemental Figure 1.**
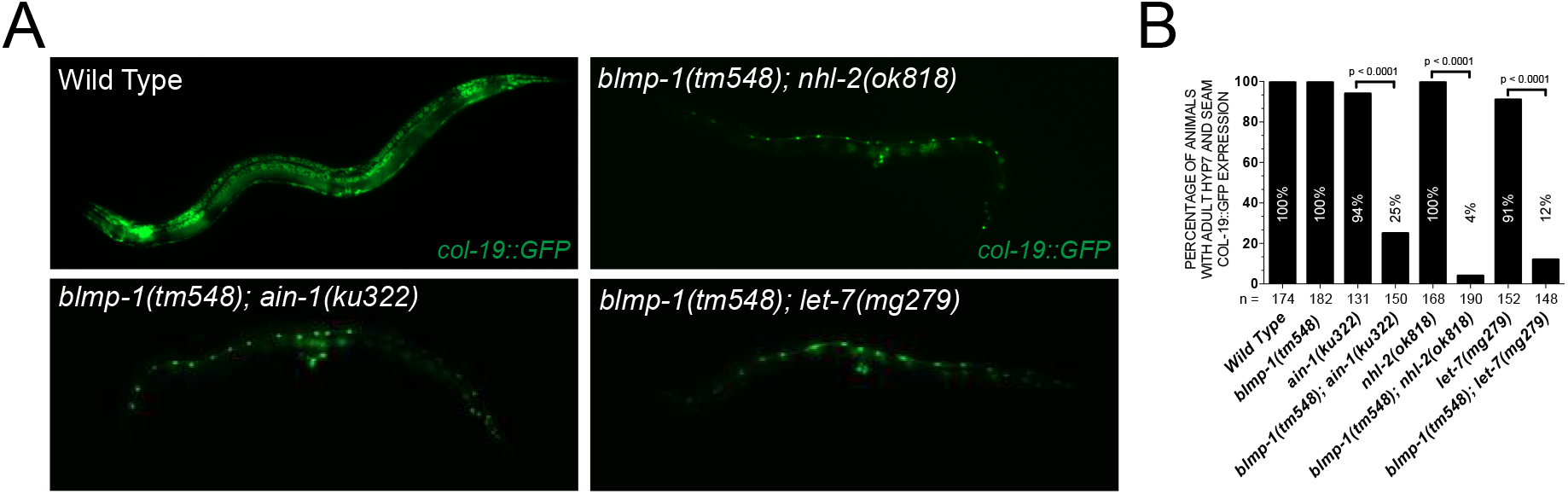
*blmp-1(0)* mutations enhance the heterochronic phenotypes other reiterative mutants. **(A)** The normal expression pattern of a *col-19::GFP* transcriptional reporter is altered in various, weak reiterative heterochronic mutants. This transcriptional reporter is usually expressed in both the lateral seam and hyp7 cells of the skin. While most mutant animals exhibit the wild-type expression pattern, combining these mutations with *blmp-1(0)* dramatically alters the expression patterns. A majority of double mutant animals only express *col-19::GFP* in the lateral seam cells. In addition, most double mutant animals inappropriately reiterate L4 seam cell division patterns resulting in supernumerary seam cells as adults. **(B)** Quantification of the *col-19::GFP* expression phenotypes in indicated mutant animals.

**Supplemental Figure 2.**
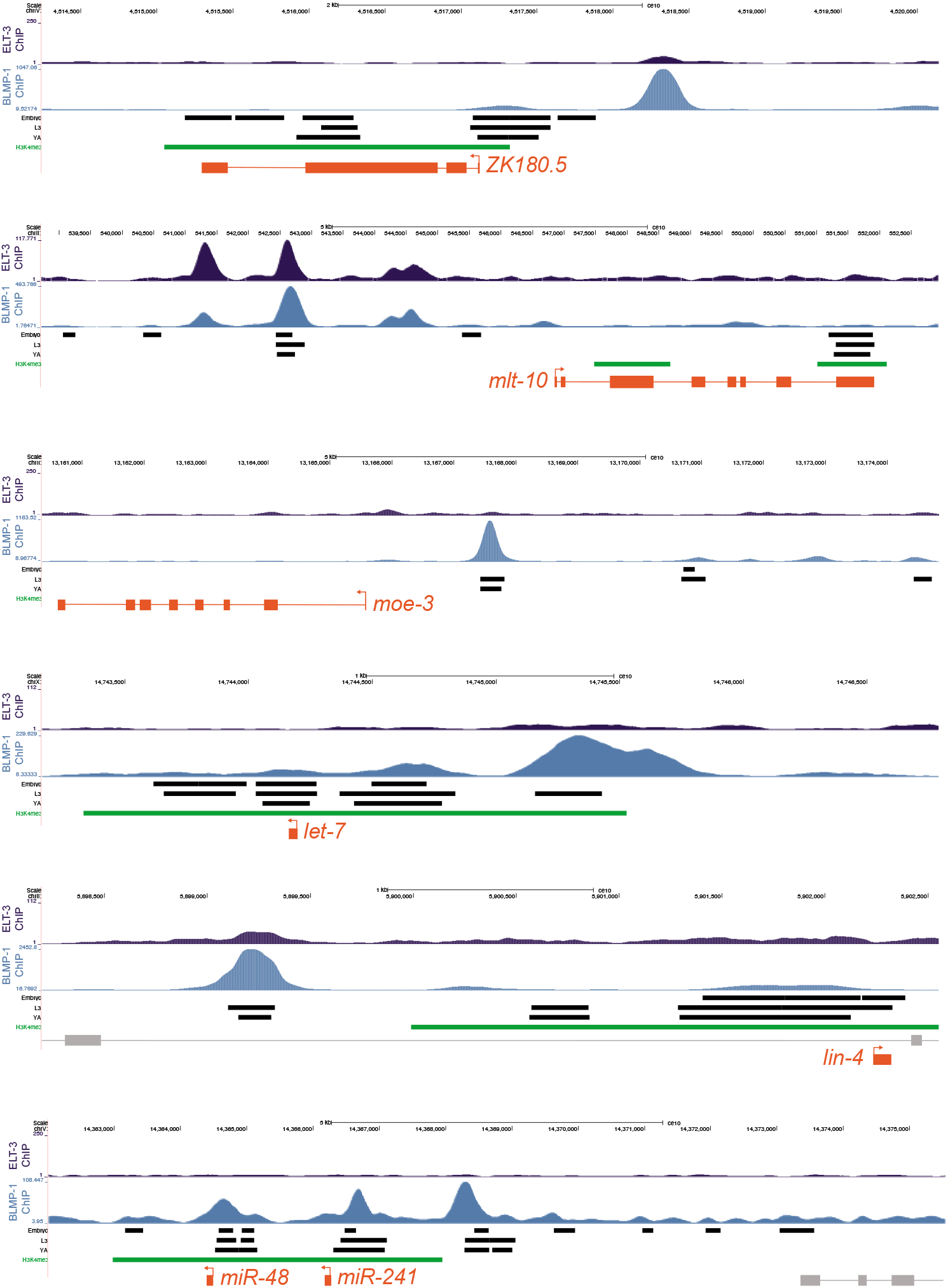
BLMP-1 and ELT-3 binding sites are enriched in the promoters of cyclically transcribed genes and are near sites of dynamically open chromatin. Browser tracks overlaying the BLMP-1::GFP and ELT-3::GFP ChIP-seq signal from L1-staged larvae for the indicated genomic loci. Green bars indicate H3K4me3 ChIP-seq data tracks (indicative of active promoters near transcriptional start sites). Black bars indicate the localization of open chromatin as measured from ATAC-seq experiments in extracts derived from embryonic, L3-staged or adult animals (Daugherty et al., 2017).

**Supplemental Figure 3.**
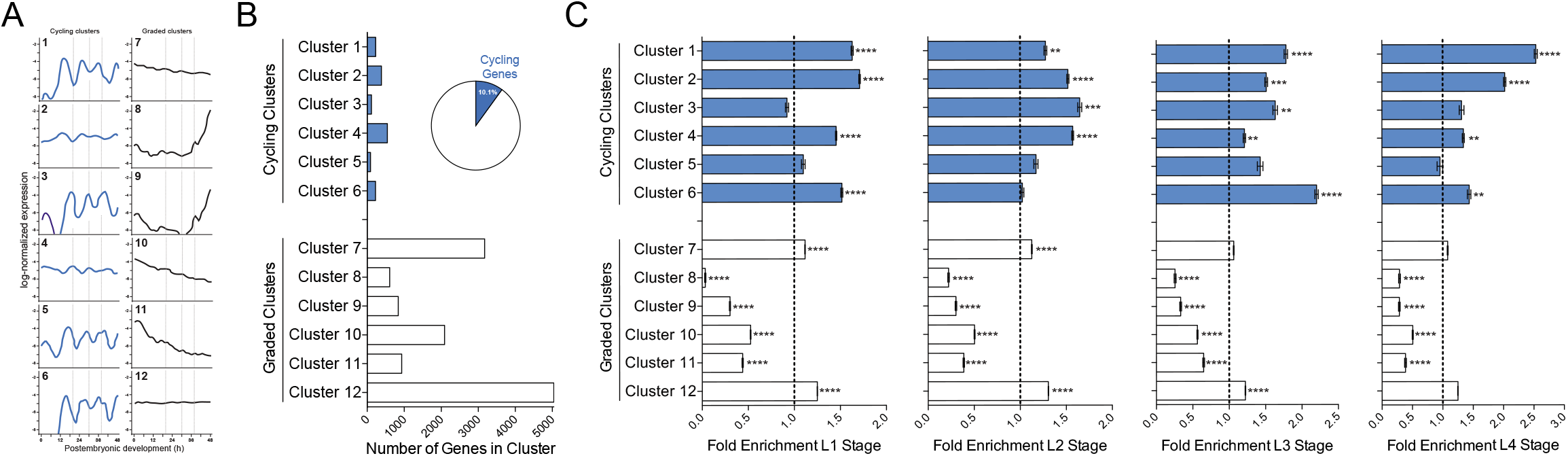
BLMP-1 binding sites are enriched in the cyclically expressed classes of transcripts identified in the Kim et al. high-resolution transcriptome data. **(A)** Clusters of genes with cycling or graded expression patters as defined by Kim et al. 2013 (Kim et al., 2013). Graphs for each cluster illustrate the average, normalized expression profiles. dashed grey lines indicate transitional between larval stages at 20°C. The cluster number is indicated in the upper left corner of each graph. **(B)** A graph illustrating the distribution of genes in each expression cluster. Pie chart illustrates the proportion of genes (10.1%) from the transcriptomic data that produce mRNA that are in cycling clusters one through six. (C) The predicted enrichment of BLMP-1 targets in each of the twelve expression clusters defined in the Kim et al. 2013 mRNA-seq analysis. Error bars are based on random counting statistics. *P* values were computed by Fisher’s exact test. Data are shown for all clusters individually. ** indicates a p values < 10^−2^ and **** indicates a p value < 10^−4^. All calculations can be found in Table S3.

**Supplemental Figure 4.**
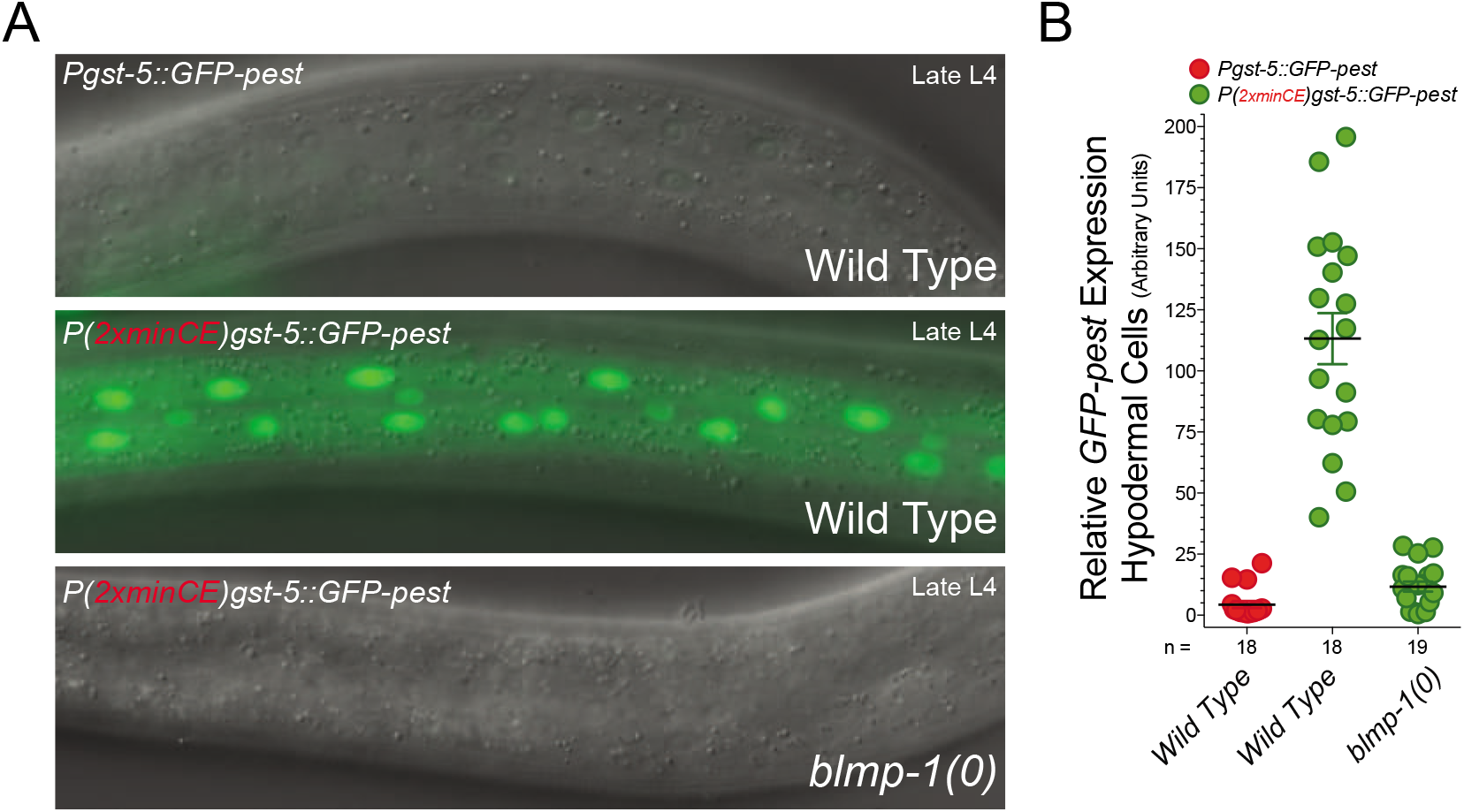
Cyclical expression of the *(2xminCE)gst-5::GFP-pest* transgene is dependent on blmp-1 activity. **(A)** Pictographs exhibiting expression of *gst-5::GFP-pest* and *(2xminCE)gst-5::GFP-pest* reporters in wild type and *blmp-1(0)* mutant animals in the mid L4 stage of development. **(B)** Quantification of individual expression levels of for the *gst-5::GFP-pest* and *(2xminCE)gst-5::GFP-pest* reporters. n = > 18 for each reporter and or genetic context. Expression of the *gst-5::GFP-pest* reporter was not altered in *blmp-1(0)* animals (data not shown).

**Supplemental Figure 5.**
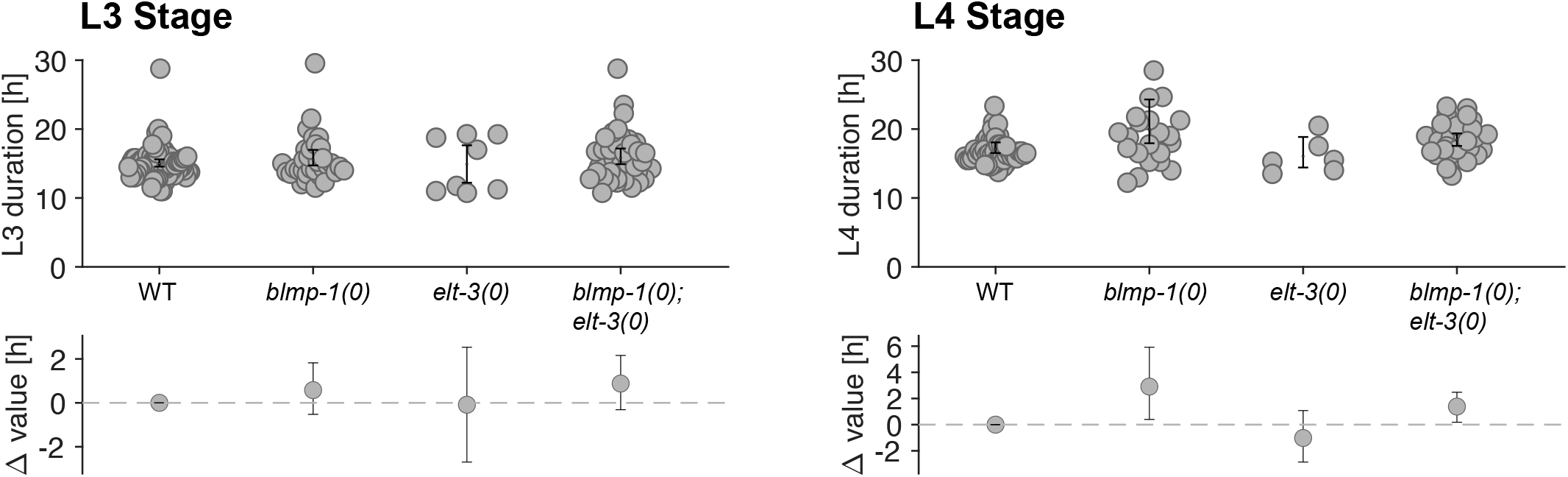
Duration times of larval stages are similar in wild-type, *blmp-1(0), elt-3(0)* and *blmp-1(0); elt-3(0)* mutant backgrounds. **(A)** Estimation plot (Ho et al., 2019) showing the duration of second larval stage (L3) for WT (N=81), *blmp-1(0)* (N=36), *elt-3(0)* (N=8) and *blmp-1(0); elt-3(0)* animals (N=40). (B) Estimation plot showing the duration of second larval stage (L4) for WT (N=51), *blmp-1(0)* (N=22), *elt-3(0)* (N=6) and *blmp-1(0); elt-3(0)* animals (N=30).

**Supplemental Figure 6.**
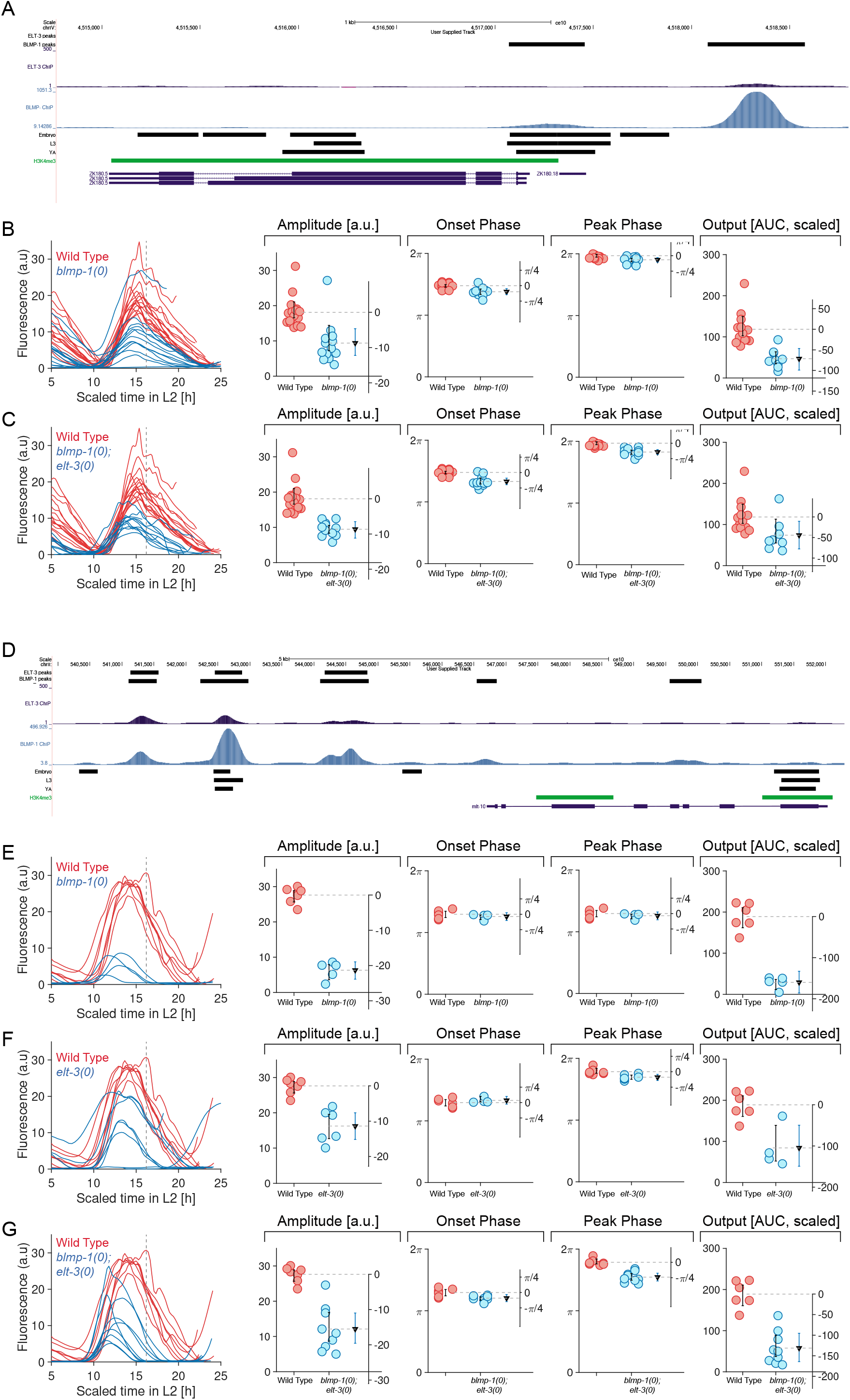
*elt-3(0)* mutations only minimally enhance expression defects of *ZK180.5::GFP-pest* and *mlt-10::GFP-pest* reporters in *blmp-1(0)* mutants. **(A)** A browser track overlaying the BLMP-1::GFP and ELT-3::GFP ChIP-seq signal near the *ZK180.5* locus. Green bars indicate H3K4me3 ChIP-seq data tracks (indicative of active promoters near transcriptional start sites). Black bars indicate the localization of open chromatin as measured from ATAC-seq experiments in extracts derived from embryonic, L3-staged or adult animals (Daugherty et al., 2017). **(B)** A comparison the L2-L3 staged traces of *ZK180.5::GFP-pest* expression and quantification of the features of these patterns in wild-type animals and *blmp-1(0)* mutants. **(C)** Same as in B except for wild-type and *blmp-1(0); elt-3(0)* mutants. **(D)** Browser track for the *mlt-10* locus illustrating the same genomic features as in panel A. **(E)** A comparison the L2-L3 staged traces of *mlt-10::GFP-pest* expression and quantification of the features of these patterns in wild-type animals and *blmp-1(0)* mutants. **(F)** A comparison the L2-L3 staged traces of *mlt-10::GFP-pest* expression and quantification of the features of these patterns in wild-type animals and *elt-3(0)* mutants. **(G)** A comparison the L2-L3 staged traces of *mlt-10::GFP-pest* expression and quantification of the features of these patterns in wild-type animals and *blmp-1(0); elt-3(0)* mutants.

**Supplemental Figure 7.**
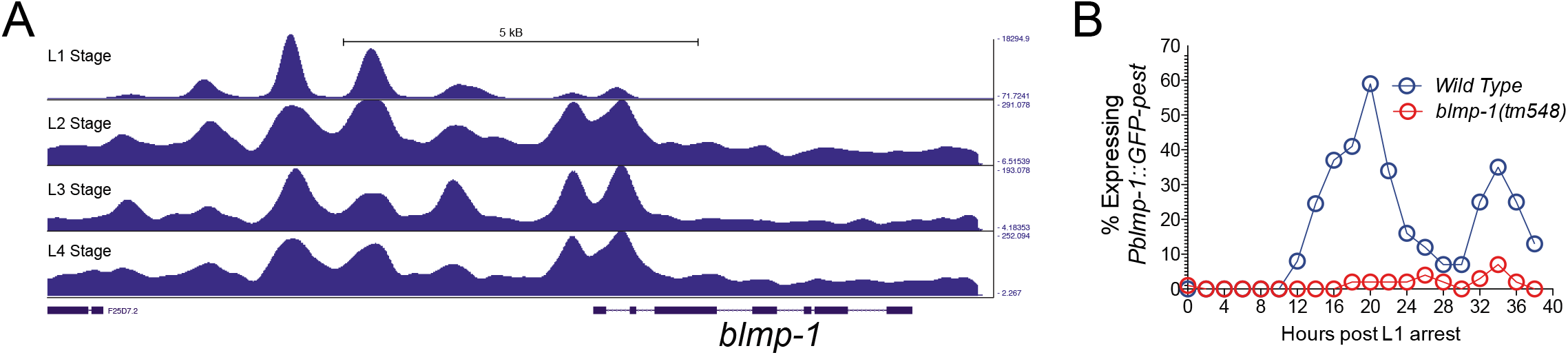
*blmp-1* expression is auto-regulatory. **(A)** Browser track of the *blmp-1* genomic region depicting the enrichment of BLMP-1 binding sites in the promoter of the *blmp-1* as defined by BLMP-1::GFP ChIP-seq signal in each stage of larval development. **(B)** A transcriptional reporter harboring a 6.2kb the upstream regulatory sequences from the *blmp-1* gene was fused to a GFP-pest reporter. Wild-type and *blmp-1(0)* animals expressing an extrachromsomal version of this transgene were stage as L1s and then plated onto normal NGM plates containing OP50. To measure the temporal expression patterns, >50 animals were scored for GFP expression every 2 hours. In wild-type animals, this reporter is expressed in a cyclical manner consistent with characterization of the endogenous *blmp-1* transcripts (Hendriks et al., 2014; Kim et al., 2013). In contrast, a majority of *blmp-1(0)* animals fail to express detectible GFP-pest expression in parallel cultures.

**Supplemental Figure 8.**
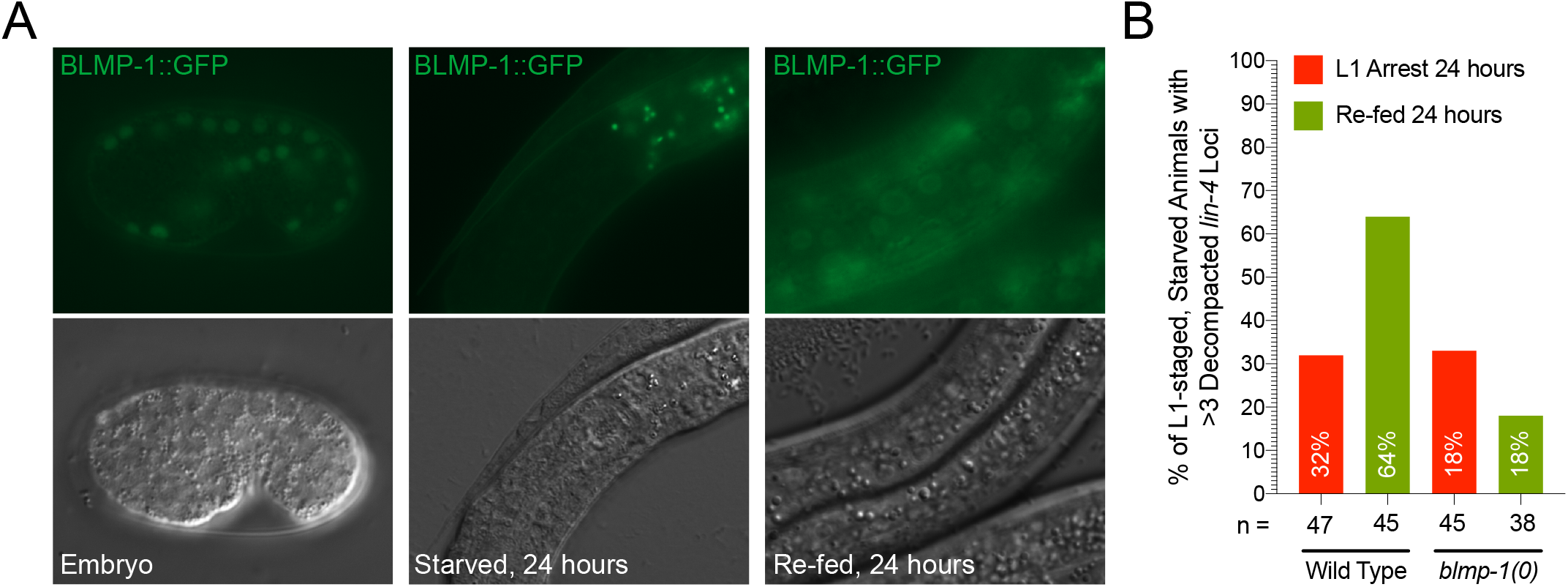
BLMP-1 expression and priming activities are repressed during L1-stage arrests. **(A)** BLMP-1 expression is present in developing embryos, reduced in arrested L1 animals and re-expressed in animals that have been re-fed. **(B)** Quantification of the number of puffed *lin-4::mCherry-pest/LacO* loci as a percentage of animals with greater than 3 puffs in the seam cells of one lateral side of animals in the indicated conditions.

## Supplemental Tables

**Supplemental Table 1. Summary of ChIP-seq experiments for BLMP-1::GFP and ELT-3::GFP.** Tables include lists of calculated peaks that were assigned to protein coding and non-coding genes for each of the 5 ChIP-seq data sets presented in Figure 3. Includes peak location, distance to assigned gene, gene name, and amplitude and phase information (if applicable). Table also includes various comparisons of ChIP-seq data sets for co-targeting.

**Supplemental Table 2. Summary of the analysis of ModEncode ChIP-seq data sets.** Includes the gene class assignments (flat, oscillating, rising; as defined by Hendriks et al.) for each TF profiled by the ModEncode project (Hendriks et al., 2014).

**Supplemental Table 3. Comparison of the enrichment of predicted BLMP-1 targets in the various expression clusters as defined by Kim et al. 2013 (Kim et al., 2013).**

